# Protein Quaternary Structures in Solution are a Mixture of Multiple forms

**DOI:** 10.1101/2022.03.30.486392

**Authors:** Shir Marciano, Debabrata Dey, Dina Listov, Sarel J Fleishman, Adar Sonn-Segev, Haydyn Mertens, Florian Busch, Yongseok Kim, Sophie R. Harvey, Vicki H. Wysocki, Gideon Schreiber

## Abstract

Over half the proteins form homo or hetero-oligomeric structures. Experimentally determined structures are often considered in determining a protein’s oligomeric state, but static structures miss the dynamic equilibrium between different quaternary forms. The problem is exacerbated in homo-oligomers, where the oligomeric states are challenging to characterize. Here, we re-evaluated the oligomeric state of 17 different bacterial proteins across a broad range of protein concentrations and solutions by native mass-spectrometry (MS), mass photometry (MP), size exclusion chromatography (SEC), and small-angle x-ray scattering (SAXS), finding that most exhibit several oligomeric states. Surprisingly, many proteins did not show mass-action driven equilibrium between the oligomeric states. For approximately half the proteins, the predicted oligomeric forms described in publicly available databases underestimated the complexity of protein quaternary structures in solution. Conversely, AlphaFold Multimer provided an accurate description of the potential multimeric states for most proteins, suggesting that it could help resolve uncertainties on the solution state of many proteins.

## Introduction

A large fraction of proteins are oligomeric in nature, often forming an assembly of multiple copies of the same folded polypeptide^1^. Functional, genetic, and physicochemical prerequisites are the driving force of the evolutionary selection of symmetrical oligomeric complexes^1–4^. The quaternary structure of a protein is of biological significance for the activity and stability of enzymes, ion channels, transcription factors, structural proteins, and more^5,6^, with the oligomeric surfaces potentially improving their stability^7^. The quaternary structure of a protein is most often determined by structural methods, such as by X-ray crystallography, which requires high protein concentration and unique buffer composition to drive crystallization. As the equilibrium between different quaternary forms is concentration and solution dependent, the registered quaternary subunit composition in the PDB or UniProt databases reflect the specific conditions applied in structure determination. While some proteins exhibit several oligomeric forms^8–10^, for most proteins only one possible assembly is indicated. A recent study showed that in *E.coli* about a quarter of proteins with known structure are monomeric, close to half are dimeric and the rest have higher oligomeric states^11^. However, that study, as well as many others, rely on the quaternary state that is indicated in UniProt or the PDB.

Several methods can quantify mass, hence quaternary structure, and shape of a macromolecular assembly. In addition to structural methods, two traditional approaches to determine oligomeric states of proteins are analytical ultracentrifugation (AUC)^12,13^ and size exclusion chromatography (SEC). The former method provides mass and stoichiometry by using sedimentation of macromolecules^14^, but requires large amounts of protein, is time-consuming, and requires expertise. The latter method, relying on the hydrodynamic radii of the proteins is straightforward but suffers from low precision. More recently, new methods have been developed that provide molecular mass with high accuracy. Here we use four different methods that can detect mass and sometimes the shape of a protein, and compare the results for 17 different *E.coli* proteins and BSA. The methods used here are native mass spectrometry (native MS), mass photometry (MP), small-angle X-ray scattering (SAXS), and SEC (alone or combined with multi-angle light scattering – MALS). All the proteins used here were produced solubly without any tag, and within a 20-35 kDa range for the monomeric state (except BSA). The abundance of these proteins in the *E.coli* cytoplasm^15,16^ covers a range from low to high levels of expression.

Native MS is a rapidly developing tool for macromolecules and protein complex investigation, maintaining the initial non-covalent interactions, hence, quaternary state, upon the transfer of solution into the gas phase at a wide range of protein concentrations ^17,18^.

MP is a new method to estimate the mass of molecules directly in solution, by quantifying their light scattering as they bind nonspecifically to a microscope cover glass^19,20^ ^21,22^. Nanogram amounts of samples are needed and it can be done in various buffers.

Small-angle X-ray scattering is an analytical method that measures the intensities of X-rays scattered by a sample as a function of the scattering angle^23^. Molecular mass (MM) is extrapolated from the forward scattering *I(0)*, using the sample concentration as input. Alternatively, the full SAXS intensities profile is fitted using the program OLIGOMER^24–26^, to produce an assembly of multimeric states for which candidate three-dimensional structures are available. In principle, information on both the average MM and shape of a protein is measured, and experiments can be conducted under various buffer conditions and across a range of protein concentrations^27^.

SEC is used for many years to analyze the oligomeric state of eluting species^28–30^. However, it is important to note that separation is based on the hydrodynamic radius of the eluting species and not the actual MM. To fix this problem, a MALS detector can be added to an SEC column, providing the absolute mass of an eluting peak^31–33^.

Here, 17 different bacterial proteins were analyzed for their quaternary structures using the four experimental solution methods detailed above and compared to reported states in PDB, SWISS-MODEL, and UniProt^34–36^. In addition, the experimental results were compared with those obtained using AlphaFold multimer^37^. For at least half the proteins studied, the oligomeric states reported in the PDB or predicted in SWISS-MODEL differed from those we identified in solution, while AlphaFold provided a much closer description to the experimentally observed oligomeric states. Native MS and SAXS measurements were performed across a range of protein concentrations, which enabled us to follow the changing equilibrium between the different oligomeric forms. While for some proteins a major shift in the predominant oligomeric form was recorded, as expected from mass action equilibrium, other proteins showed multiple, concentration independent oligomeric states, which relation could however be affected by unfolding/refolding the protein, suggesting a high transition between the different forms. To our knowledge, this is the most systematic study of oligomeric forms of proteins in solution, suggesting that our current perspective of defined oligomeric states is under estimating the real structural complexity of the quaternary structures of proteins.

## Results

### A benchmark for studying the oligomeric state of proteins

Most of our knowledge of the quaternary structure of proteins stems from structural methods, mainly X-ray crystallography, where the protein of interest is typically at a fixed, high concentration. Recent method developments provide the opportunity to revisit this question using a range of approaches. We analyzed the oligomeric state of 17 different bacterial proteins and BSA by native MS, MP, SAXS, and SEC. The results were compared to those previously reported and deposited in the following biological databases: PDB, UniProt, and PISA and to the predicted results of SWISS-MODEL^34–36^. The experimental methods used here can determine the mass and/or low-resolution structure/shape of a protein in a solution. This allowed us to evaluate the limitations and sensitivities of the different techniques.

Native MS analysis was conducted across a range of concentrations (40 - 0.15 μM, prior to the dilution inherent to the measurement), in an automated, high throughput manner. As the oligomeric composition is determined at multiple concentration points, the concentration dependent equilibrium is recorded for each sample^38^.

MP was performed at a single, low concentration (~100 nM), due to instrument limitations. MP is limited to the detection of macromolecules above a size threshold of ~40 kDa, thus the monomeric states of the proteins in our work were mostly not detected. However, higher oligomers were readily identified.

We used SAXS at four concentration points to characterize the molecular mass (MM) and to calculate a low-resolution structure of the protein assemblies. From the forward scattering intensity, *I(0)*, MM values were calculated directly from the experimental data (Table S1). In an indirect approach, using both experimental data and the high-resolution crystallographic structures, linear combinations of the computed scattering patterns from the input structures were determined that best fit the SAXS data in the program OLIGOMER^24^. Protein concentrations ranged between 10-90 μM, which enabled characterization of concentration dependent equilibria, and provided structural models of the protein assemblies present in solution.

For SEC analysis, we first calibrated the column to obtain a relation between the MM of known proteins and the elution volume (Fig. S1). The oligomeric state was calculated as the quotient of the experimentally determined MM and the monomeric MM. For cases where the quotient is not an integer, the protein may exist in dynamic equilibrium between several oligomeric forms. In the following section, we provide detailed results on the quaternary structure of a set of proteins obtained from four independent methods, from the simplest case to the more complex.

Bovine Serum Albumin (BSA) is a blood protein often used as a standard in many biophysical techniques. BSA is well characterized and is typically found in both monomeric and dimeric forms^39^. We characterized BSA at a range of concentrations using the various methods outlined above. Fig. S2 and Fig. S3 upper panel show that under the conditions used in this study, over 90% of BSA is monomeric, independent of the method and the protein concentration used. Thus, the two dominant oligomeric states observed for BSA are stable, and not connected through mass action. This observation is in line with previous publications, where a constant, minor population of dimeric BSA was typically observed^40–42^. The type I interferon protein (IFNα2) provides an example of a protein that can form a concentration dependent dimer connected through mass action. Using SEC we show in Fig. S3 lower panel how the SEC elution volume of IFNα2 is reduced at higher protein-concentrations, suggesting a higher oligomeric state at higher protein concentration. Indeed, while IFNα2 is a monomer in dilute solutions, it was solved as a zinc mediated dimer by x-ray crystallography^43^.

To obtain a more general view of the oligomerization state of proteins in solution, we analyzed the quaternary structures of 17 different bacterial proteins using the four methods described above. The proteins were expressed and purified with a cleavable His-tag and a sumo protease cleavage site as described in^44,45^. This allows proteolysis and elution directly from the Ni-NTA beads, without leaving a trace of the linker protein (see materials and method). Fig. S4 shows SDS-Page analysis of the 17 proteins used in this study, with and without reducing agent. With the exception of IspD, all proteins run primarily as monomers also in the absence of β-mercaptoethanol, showing that covalent inter-protein disulfide bridges are not present in the purified proteins.

### SodA is a classic example for a dimeric protein

Superoxide dismutase (SodA), a 23 kDa protein responsible for the destruction of toxic superoxide radicals within living cells^46^. In the PDB, UniProt, SWISS-MODEL, and PISA, SodA is described as a homodimer. Native MS analysis performed at concentrations of 0.94 - 40 μM showed SodA to be a dimer (Fig. 1A and B). MP analysis showed a single peak at 52 kDa (Fig. 1D), which we assume is also the dimer. The SEC elution volume of SodA (Fig.1C) corresponds to a molecular weight of 36 kDa (28-45 kDa), which suggests a monomer-dimer equilibrium. While the forward scattering intensity of the SAXS data (*I(0)*) for SodA suggests a dimeric species across all measured concentrations (Table S1), a fit of the dimeric crystal structure (PDB id: 1D5N) provides a less satisfactory description of the solution data than that derived from the equilibrium fitting procedure (Fig. 1F). Optimal fits to the scattering data at each concentration point are observed when the data are described as a concentration dependent monomer and dimer equilibrium, with a very small proportion of SodA tetramer (a dimer of dimers was extracted from the crystallographic unit cell, Fig. 1E–F). As the concentration is increased, the equilibrium shifts toward the dimeric state, and it is expected that structural methods, using high protein concentrations favor the formation of SodA dimers. The SodA assemblies used for the equilibrium description of the data are shown in Fig. 1G.

**Figure 1:**
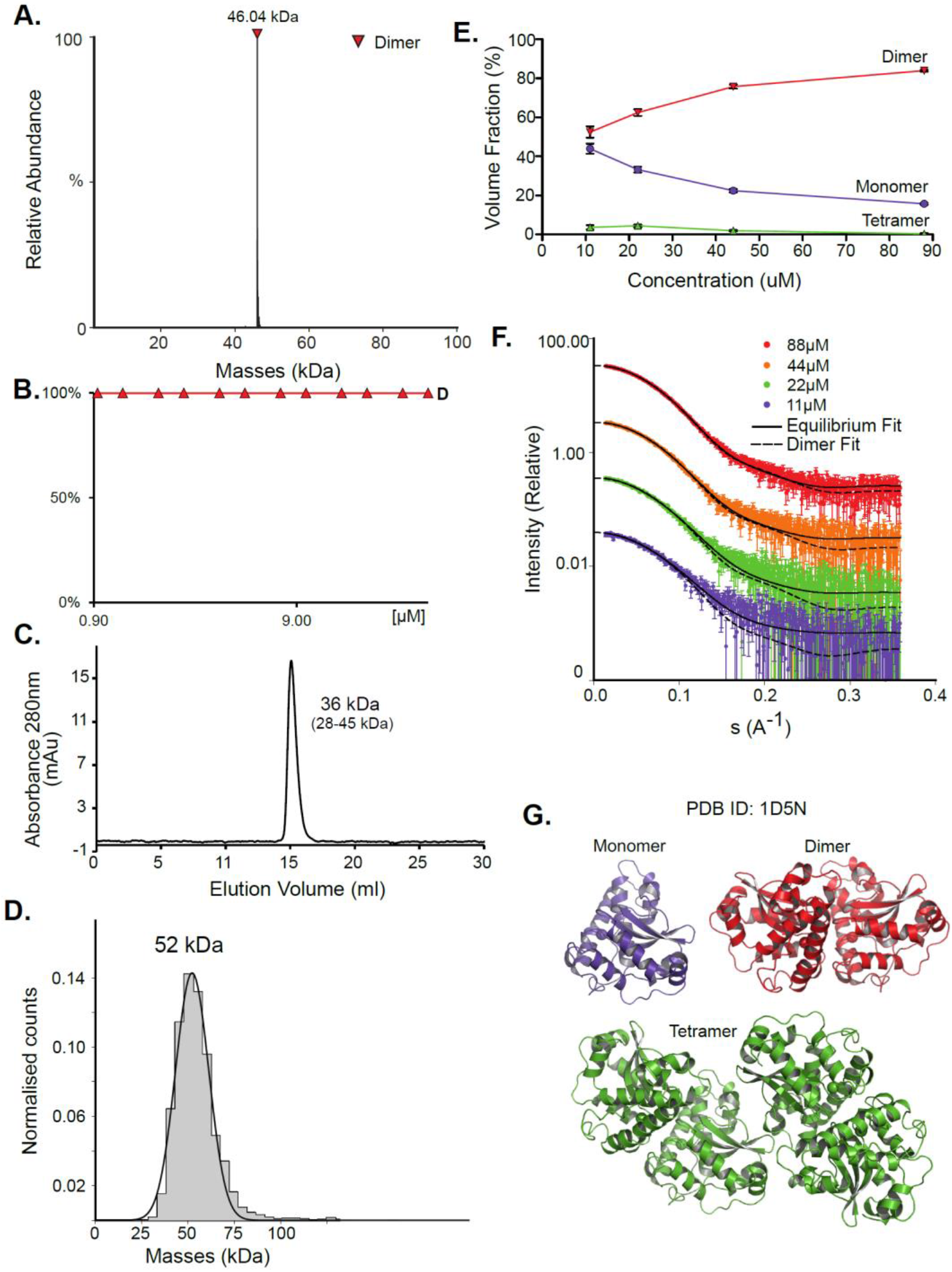
SodA oligomeric composition as determined by multiple methods. **A.** Native MS returns a single peak, corresponding to the mass of a dimer of 46.04 kDa at all concentrations. UniProt MM of a monomer- 22.97 kDa. **B.** Native MS in a range of protein concentrations, from 0.9μM to 40μM, shows a stable dimer. **C.** SEC analysis shows one main peak that eluted at 15.08 ml, corresponding to 36 kDa (28-46 kDa). This would suggest over one but under two subunits by the MM calculations (see Fig. S1). **D.** Mass photometry measurements of the protein shows a mass corresponding to the dimeric state. **E.** SAXS equilibrium fitting using the program OLIGOMER and PDB id: 1D5N suggests an equilibrium between a monomer and a dimer, with the volume fraction of dimer increasing with concentration. A small almost constant population of a tetramer is also present, solely seen in this method at low concentration (10-20μM). **F.** Raw SAXS data fitted by OLIGOMER at four different protein concentrations, 11 μM (purple), 22 μM (green), 44 μM (orange), and 88 μM (red). Fitted lines are represented by the black line. Fitting the data to only a dimeric state is shown in dashed black lines. **G**. Input assemblies of SodA, PDB id: 1D5N, used for SAXS equilibrium analysis.

### DeoC shows a mass action driven monomer/dimer equilibrium

While for SodA most methods predict a dimer, the picture for Deoxyribose-phosphate aldolase (DeoC) is more complex. DeoC is a protein with a monomeric molecular weight of 27.7 kDa, responsible for the catalysis of a reversible aldol reaction between acetaldehyde and D-glyceraldehyde 3-phosphate, to generate 2-deoxy-D-ribose 5-phosphate^19,20^. The structure of this protein was solved independently in both dimeric (PDB:1KTN) and monomeric (PDB: 1JCL) states. In UniProt (P0A6L0) it is ambiguously designated as both a monomer and dimer^9,47^. Using native MS, we observe a concentration dependent monomer-dimer equilibrium (Fig. 2A–B). Two peaks are observed by native MS, with the relative abundance of the peaks being strongly affected by the protein concentration. At lower concentrations the monomeric form is dominant whereas in higher concentrations the dimer is predominant (Fig. 2A). The equilibrium dynamics are shown in Fig. 2B, where a shift in oligomeric composition clearly follows a change in concentration. Fitting the data to eq. 3:

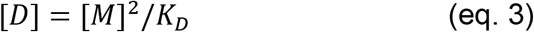

where D is the dimer and M monomer gives an apparent affinity of 2.2 μM for the monomer-dimer equilibrium (Fig. 2C). SAXS measurements modeled well the monomer-dimer equilibrium shown by native MS (Fig. 2G–H). As the SAXS was done only at higher concentrations, a full binding curve could not be constructed but the equilibrium follows nicely the trend described by the native MS data. SAXS MM values calculated from *I(0)* values show it to be a dimer at all concentrations (Table S1). SAXS also provides structural models for both the monomer and dimer forms (Fig. 2G–H). SEC measurements show a single elution peak that corresponds to 45 kDa (35-57 kDa) which is between a monomer and a dimer, closer to a dimer (Fig. 2D). DeoC was injected at a concentration of 21.6 μM, which is diluted during its progression in the column. Mass photometry measurements show two peaks, 36 kDa and 55 kDa (Fig. 2E). The first is probably the tail of the monomer, as the method has a 40 kDa detection threshold. Therefore, we are unable to determine the ratio between the peaks, and the exact monomeric mass. However, the molecular weight of the dimer matches the expected mass of 55 kDa. In MP we applied a concentration of 53 nM, which is lower than used in the other methods. Therefore, we assume the monomeric peak to be much larger than observed. Concentration dependence of the oligomeric state is a direct outcome of mass action behavior of molecules and has been previously reported for other proteins^48^ as shown above for IFNɑ2 and BSA.

**Figure 2:**
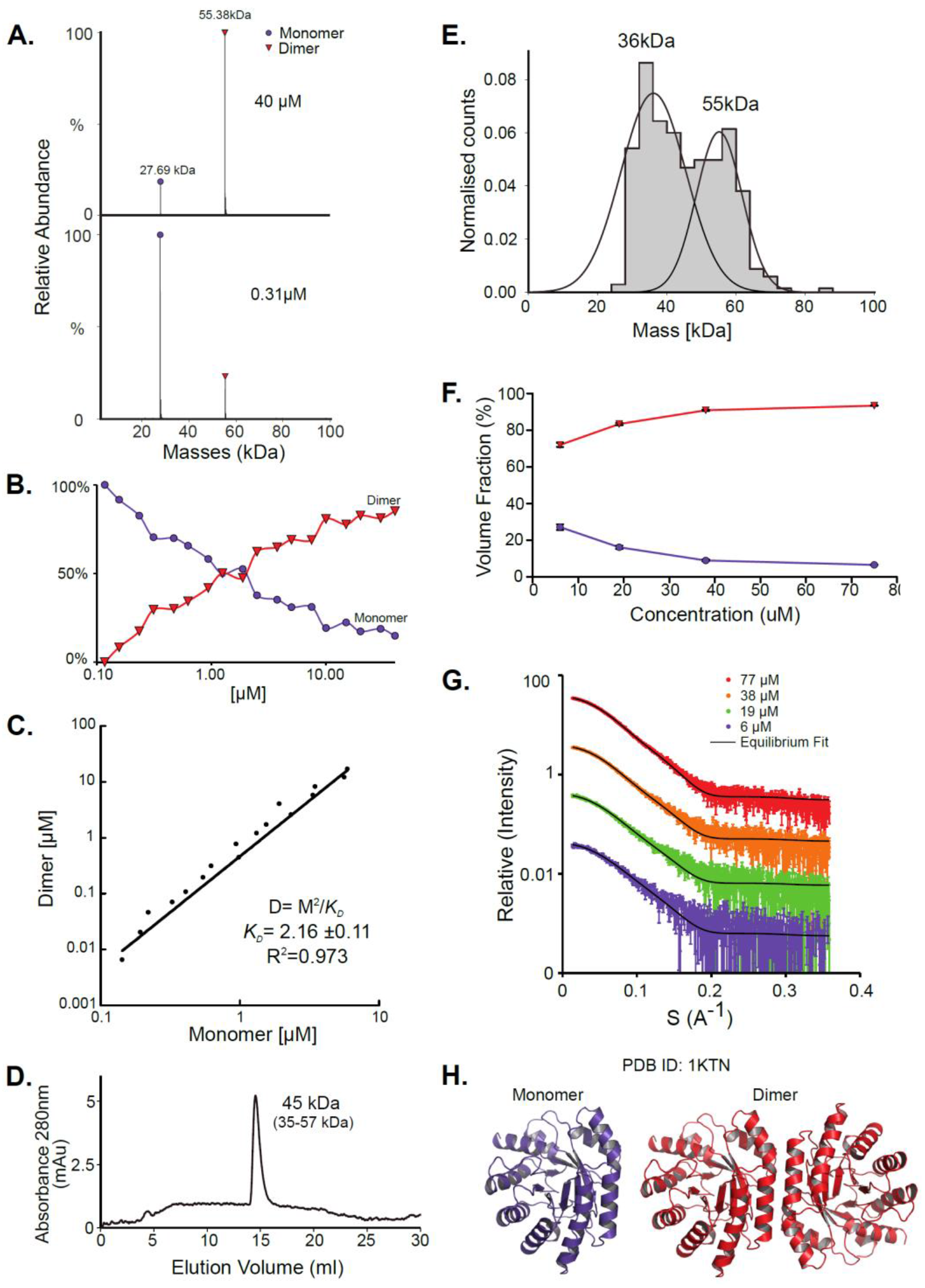
DeoC exist in a concentration-dependent monomer/dimer equilibrium. **A.** Native MS results show two peaks that correspond to a monomer and a dimer with MM of 27.69 kDa and 55.38 kDa respectively. UniProt MM of monomeric DeoC- 27.74 kDa. **B.** Molecular mass as determined by native MS in a range of protein concentrations, from 0.12 to 40 μM. **C.** Fitting the monomer-dimer equilibrium using equation 3 (M-monomer and D-dimer concentrations). **D.** SEC analysis shows one main peak that eluted in 14.52 ml, corresponding to 45 kDa (35-57 kDa), which is between a monomer and a dimer. **E.** Mass photometry measurements of the protein are showing a mass that fits monomeric and dimeric states of the protein, the MP mass threshold is around 40 kDa so the monomeric observed peak is probably the tail of a much larger, not observed peak. **F.** SAXS data were fitted using the program OLIGOMER with PDB id: 1KTN for modeling the structures. At the concentrations used, most of the protein is in dimeric form. **G.** Equilibrium fit of SAXS data at four different protein concentrations, 6 μM (purple), 19 μM (green), 38 μM (orange), and 77 μM (red). Fitted lines are represented by the black line. **H**. Assemblies of DeoC, PDB id: 1KTN, used for SAXS equilibrium fitting.

### FabG is in multiple concentrationindependent oligomeric states

FabG, (3-oxoacyl-[acyl-carrier-protein] reductase), catalyzes the NADPH-dependent reduction of β-ketoacyl-ACP substrates to β-hydroxyacyl-ACP products, the first reductive step in the elongation cycle of fatty acid biosynthesis^49–51^. The monomeric unit of FabG has a MM of 25 kDa, while the structure of the *E.coli* FabG protein shows it to be a tetramer. An analysis by the Protein Quaternary Structure Investigation database PiQsi,^52^ that provides manually annotated sizes of biological units from the literature for PDB entries, predicts FabG to be a tetramer. In UniProt (P0AEK2) its subunit structure is registered as homo-tetramer. Therefore, we were very much surprised to find this protein to be mostly hexameric in solution, with a smaller percentage of dimer and tetramer (Fig. 3). The native MS data shows that the oligomerization state of the protein is predominantly hexameric, independent of the concentration used in this study (Fig. 3A–3B). SEC analysis shows one main peak and an additional smaller peak, corresponding to ~101 kDa and 41 kDa species (tetramer and a dimer). Strangely, the hexameric state is not seen here, perhaps due to a hydrodynamic radius similar to that of a tetramer, and thus poor peak resolution of such species (Fig. 3C). The MP analysis shows a very similar picture to native MS with dimeric, tetrameric and hexameric forms observed (Fig. 3D). As FabG was diluted from 112 μM to 38 nM before the MP measurements, we could investigate time dependent changes in the oligomeric state after dilution. Measuring the oligomeric state at four time points after dilution (from immediately after dilution to overnight, Fig.S5A), shows the fractions of the different oligomers to be static with time, with the hexamer and dimer being the major species. SAXS data for FabG was fitted by OLIGOMER using the high-resolution crystallographic structure (PDB: 1I01), and could not be satisfactorily described by the tetrameric form and dissociation products. A hexameric rigid body model was generated from the 1I01 monomer subunit using the program SASREF and with P32 symmetry. This rigid body model, in combination with the 1I01 dimeric and monomeric assemblies provided a very good fit to the experimental data. In line with the MS and MP results the SAXS equilibrium analysis also identifies the hexamer to be the dominant species with a minor population of monomer at low concentration (Fig. 3F–H). The SAXS MM as calculated from *I(0)* values is close to that of a tetramer (3.5 times the mass of a monomer), however, as the tetrameric structure does not fit the experimental SAXS curve, *I(0)* is assumed to describe an equilibrium of multiple-oligomeric states (Table S1). To validate whether differences in protein-refolding had an effect on the oligomeric states in solution, we unfolded FabG in 8M Urea before buffer exchange back to PBS. Figure 3E shows the oligomerization state after refolding, as determined by MP. Again, a mixture of oligomeric states was observed, but now the dimer and tetramer were the most abundant, with a smaller fraction of hexamer. Changing the pH and salt concentrations (Figures S5B-C) had no drastic effect on the observed oligomeric forms of FabG.

**Figure 3:**
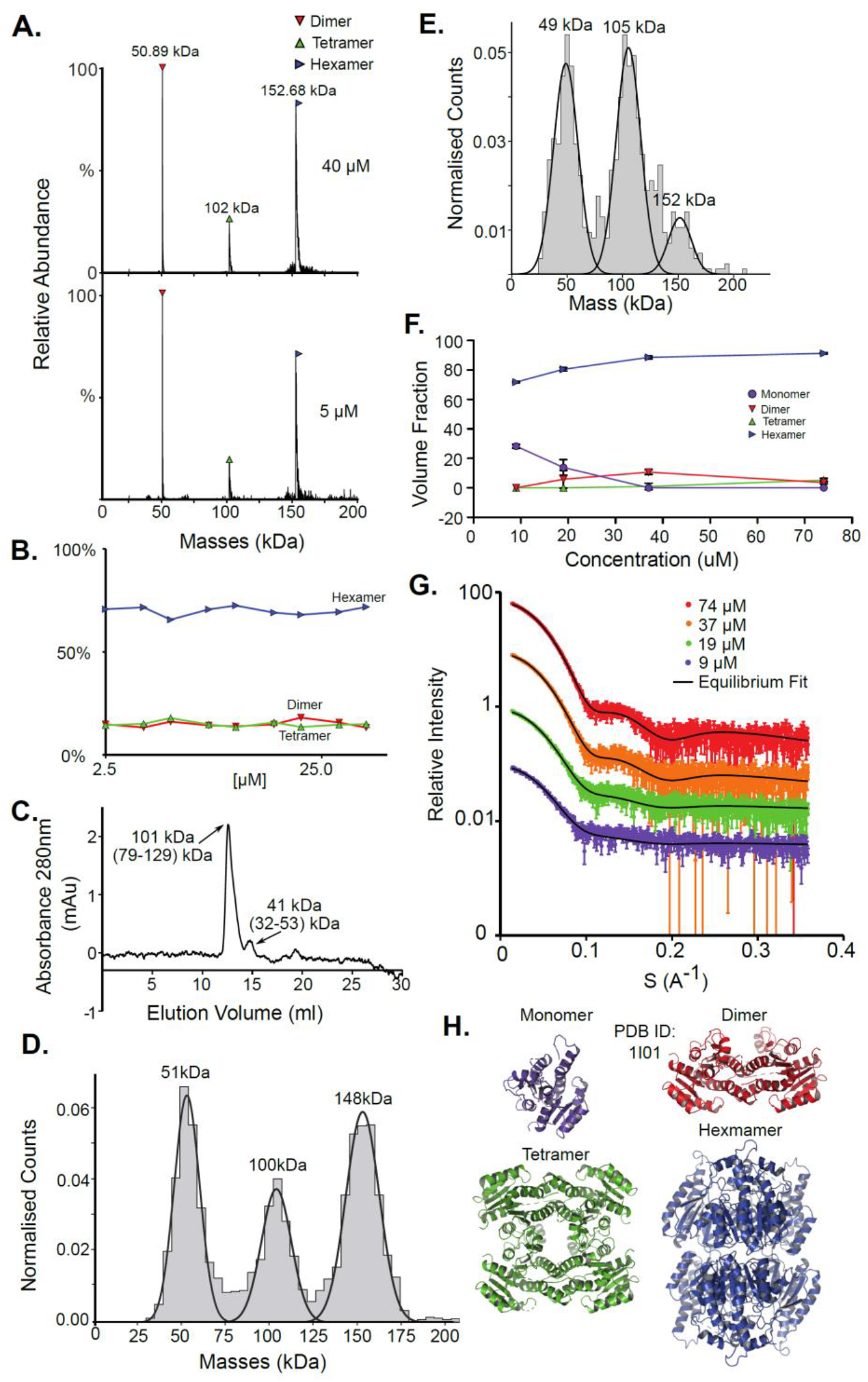
FabG is in dimer, tetramer, hexamer equilibrium. **A.** Native MS results show three peaks that correspond to a dimer, a tetramer, and a hexamer, 50.9 kDa, 102 kDa, and 152.7 kDa accordingly. UniProt MM of a monomer- 25.56 kDa. **B.** Native MS in a range of protein concentrations, 2.5 μM - 40 μM, shows the proteins oligomeric state to be independent on the concentration. **C.** SEC analysis shows one main peak eluted in 12.56 ml, corresponds to 101 kDa (tetramer) and a smaller peak that elutes at 14.72 ml (41 kDa) corresponding to a dimer by mass calculations (see Fig.S1). **D.** MP measurements of the protein shows masses that fit a dimer, a tetramer and a hexameric state. **E.** Mass photometry measurements of FabG after unfolding in 8M urea for 30 minutes and refolding back to PBS shows a reduction in occupancy of the hexamer. **F**. SAXS equilibrium fitting using the program OLIGOMER and PDB id: 1I01, shows mostly hexamers, with some dimers and monomers at lower concentrations. A small population of a tetrameric state, is also observed. **F.** Equilibrium fit of SAXS measurement in four different protein concentrations, 9 μM (purple), 19 μM (green), 37 μM (orange), and 74 μM (red). Fitted lines are represented by the black line. **G**. Input assemblies of FabG, PDB id: 1I01, used for SAXS equilibrium fitting.

### NadK is in multiple concentrationdependent oligomeric states

NadK is a key enzyme in the biosynthesis of NADP^+^, catalyzing the phosphorylation on 2'-hydroxyl of the adenosine moiety of NAD^+^ to yield NADP^+^ ^53,54^. Its monomeric MM is 32.5 kDa. While the structure of *E.coli* NadK was not solved, the structure of NadK from *Yersinia pestis* (82.5% sequence similarity) has been determined (PDB: 4HAO), from which a homology model was built using SWISS-MODEL. The structure predicts NadK to be a dimer, however, PDB 4HAO suggests it to be a tetramer. Indeed, native MS detected a dimer-tetramer equilibrium, with trace amounts of monomers observed (Fig. 4A and B). The ratios between these three species are concentration dependent. At low concentrations (~1 μM) the percentages of the different species were 12% monomer, 56% dimer, and 32% tetramer. At high concentrations (~40 μM), the ratio between the species shifted to 2% monomer, 23% dimer and 75% tetramer. SEC analysis shows one main peak that corresponds to 88 kDa, which is between 2 to 4 subunits (Fig 4C). This would reflect a dynamic equilibrium between dimer and tetramer in solution, which might reflect the ratio of the two in the solution. NadK was measured by MP at a concentration of 88 nM, showing a similar oligomerization pattern as observed by native MS at low protein concentrations, with the dimer being the more dominant form. However, small fractions of additional hexameric and octameric states were also observed (Fig. 4D). The SAXS-determined MM from *I(0)* was in line with the average native MS data at similar concentrations and consistent with that for an equilibrium of oligomeric states, yielding MM of 77 kDa at 8 μM and 87 kDa at 61 μM protein concentrations (corresponding to subunit averages of 2.37 and 2.67, respectively) (Table S1). However, using the fitting procedures of the program OLIGOMER and the *Yersinia pestis* structure, an equilibrium between monomer, tetramer and octamer was identified, with no dimer detected. MM estimates independent of the concentration values were calculated from the hydrated particle volume (Porod volume) extracted from the SAXS data. These suggest an average MM much higher than that of a tetramer at the highest concentrations measured by SAXS. The NadK data are well described by an equilibrium mixture with the main component being a tetrameric arrangement (defined as biological assembly 3 by the authors of the 4HAO crystal structure), a small amount of monomer, and an aggregate modeled as a dimer of tetramers (8-mer) (Fig. 4G).

**Figure 4:**
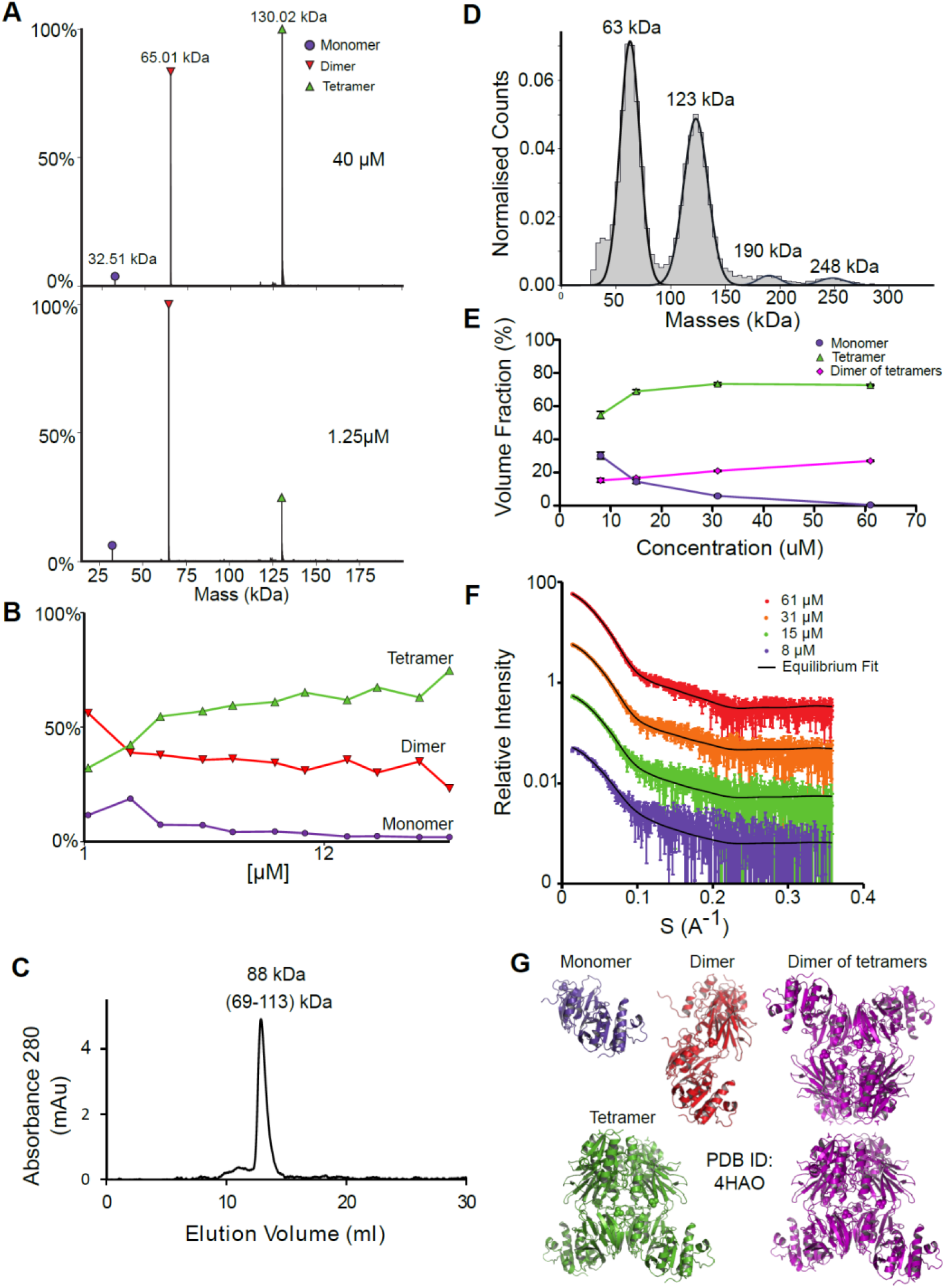
NadK oligomerization equilibrium. **A.** Native MS results show three quaternary states of the protein; a monomer, a dimer, and a tetramer. UniProt MM of a monomer: 32.57 kDa. **B.** Native MS of NadK in a range of protein concentrations, 1.25μM - 40μM. At 40μM, 75% of the protein is in a tetrameric state, 23% is a dimer and only 2% are monomers. At 1.25 μM the dimeric state is the most populated one, followed by tetramer and monomer. **C.** SEC analysis shows one main peak that eluted in 12.9 ml, corresponding to 88 kDa (2.7 subunits by the mass calculations - see Fig. S1). **D.** MP measurements of NadK revealed four masses that fit its multimeric states: dimer, tetramer, hexamer, and an octamer. The highest two multimers are found in low percentages. **E.** SAXS equilibrium fitting using the program OLIGOMER and PDB id: 4HAO shows an equilibrium of predominantly tetrameric NadK, with a concentration-dependent fraction of monomer and a larger assembly, described here as a dimer of tetramers. **F.** Equilibrium fit of SAXS measurement in four different protein concentrations, 8 μM (purple), 15 μM (green), 31 μM (orange), and 61 μM (red). Fitted lines are represented by the black line. **G**. Input assemblies of NadK used for the SAXS equilibrium fitting. Note that PDB id: 4HAO used here corresponds to the homologous *Yersinia pestis CO92* protein with 82.5% sequence similarity to the *E.Coli* NadK.

In addition to the five proteins presented here in detail, we measured the multimerization state of 12 additional proteins using most of the methods described above (Table S2). An example of another protein where a dynamic equilibrium was observed is Upp, which is mostly a dimer, but with the fraction of tetramer and hexamer increasing at higher protein concentrations. Diluting concentrated Upp at time 0 and following its multimerization state with time shows a fast transition between these different states (Fig. S6), with the MP measurement done about one minute after dilution showing the different multimeric states, while 30 minutes later the protein is only a dimer. This experiment clearly shows that Upp is in a concentration-dependent equilibrium between different oligomeric states, as also shown in Table S2 and Fig. S6.

### AlphaFold Multimer predicts potential oligomeric states

AlphaFold2 (AF) has demonstrated atomic-level accuracy in *ab initio* prediction of the structures of monomers^55^ and protein complexes^37^. Here, we examined whether AF can also predict the oligomeric state of homo-oligomers by comparing its predictions to our experimental results. Prediction quality was evaluated based on the AF confidence parameters ipTM (overall predicted model accuracy weighted more heavily at the oligomeric interfaces) and PAE (confidence in the relative orientation of the monomers). When ipTM scores are low (<85%), visual inspection often reveals backbone clashes, deviations in the monomer compared to available PDB structures, or unrealistic intermonomer orientations. By contrast, predictions with ipTM > 85% exhibit accurate monomer structures and symmetric intermonomer interactions.

AF always predicts the monomeric state, even when this state is not detected experimentally at the given concentrations. In 13 out of 17 cases, AF correctly predicts the prevalent multimeric state, including ThiD for which no experimentally determined structure is available (Fig. 5A). In 12 of these, the monomer structures exhibit low RMSD to available PDB structures (< 1.1 Å). For Can and Upp, AF assigns ipTM scores >85% to both the tetramer and the dimer. Visual inspection reveals that the dimer corresponds to half of the tetrameric state (Fig. 5B). In the case of CAN, AF also predicts a trimer that was not observed experimentally. Visual inspection of the predicted trimer shows an obvious gap in symmetry, such that this trimer is a subset of the tetrameric model (Fig. 5C). For the hexameric SpeB, AF assigns a marginal ipTM score of 83% to the trimer form. Visual inspection reveals that the predicted trimer is half the hexamer. In the specific case of FabG, for which the hexamer is one of the prevalent forms (Fig. 3), the predicted model lacks confidence (ipTM 76%). Nevertheless, the AF hexameric model shows accurate monomeric structures for each monomer (RMSD to PDB entry 1I01 <1.4 Å). The hexamer also exhibits C6 symmetry and only subtle side chain clashes that are readily relieved using Rosetta whole-protein minimization and yields a low energy structure (Fig 5D). The low AF confidence, however, means that this prediction may only serve as a working model that requires further substantiation. In the case of the IspD, the experiments show a prevalent dimer and a minor tetrameric species. AF provides a confident dimer prediction whereas the tetrameric model receives low scores and reveals substantial backbone clashes. In only two cases, NfuA and BaeR, AF is unable to recapitulate the monomeric structure, and subsequently, no high-scoring multimers are predicted. Both of these proteins comprise two domains connected by a long disordered region. Based on plDTT, PAE scores and visual inspection, the domains are predicted to be folded, but the orientation between them is uncertain. In the case of NfuA, one of the domains exhibits a structure that is close (<1 Å RMSD) to the structure of an orthologue from *Arabidopsis thaliana* (PDB code: 2z51; sequence identity 38%), and in the case of BaeR, both domains are predicted with RMSD < 0.6 (PDB code: 4b09) but their orientation is incorrect. Subsequently, all higher multimers of these proteins resulted in backbone clashes, and low ipTM scores <50%.

**Figure 5.**
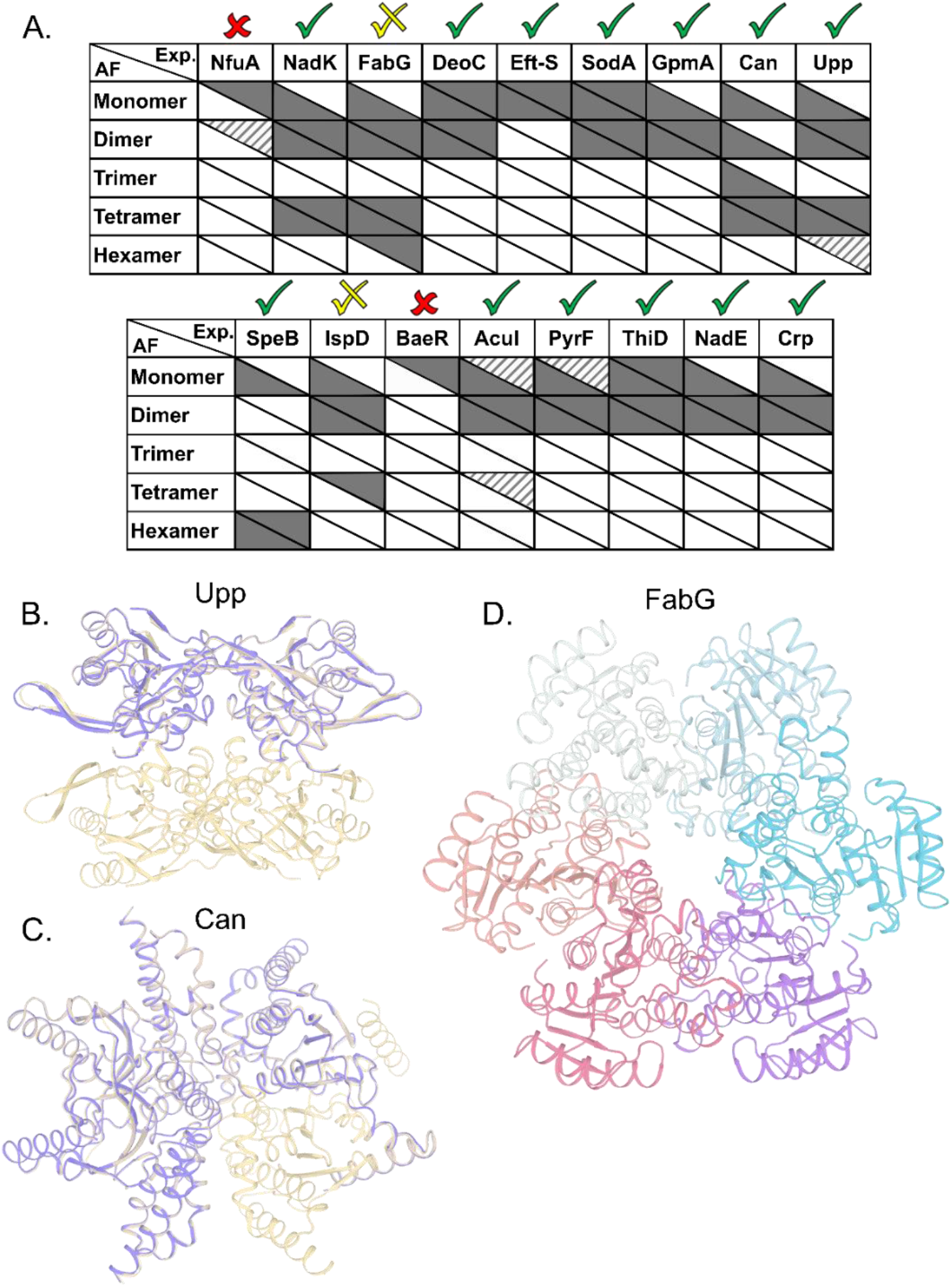
Correspondence between experimentally observed homo-oligomeric states and AF predictions. A. Upper triangle of each cell indicates the prevalence of the multimeric states found experimentally, with solid, striped and no color representing high, low and unobserved species respectively. The lower triangle of each cell shows AF’s prediction accuracy, with solid color cells representing models with ipTM scores above 85%, and no color is ipTM below 85%. For monomers, mean lDDT scores were used instead of ipTM, with >90% as solid color, 80%-90% stripped, and <80% white. Checks above columns indicate the usefulness of the prediction in determining the main multimeric state assed by us based on ipTM, PAE scores and visual inspection of the structures. B. Upp structure prediction of a dimer (purple) and tetramer (wheat). C. Can structure prediction of a trimer (purple) as a subset of the experimentally validated tetramer form (wheat). D. AF prediction for FabG in hexameric form after Rosetta relaxation.

In summary, AF Multimer was able to predict well the multimeric forms of the different proteins compared to empirical methods tested here. Predictions were better for dimers and tetramers, while hexamers are still difficult to predict using AF.

## Discussion

Quaternary composition is an important factor in the overall assembly of proteins. In contrast to the primary, secondary, and tertiary structure, the quaternary assembly is made of non-covalently interacting subunits, and thus their assembly is a higher order reaction, which also depends on the protein concentration. Therefore, a description of a protein as monomer/dimer/tetramer etc. is a simplification, which does not consider the conditions under which this assembly was determined. Moreover, from the law of mass action, we expect protein quaternary structures to be in an equilibrium between multiple forms, if specific forms are not kinetically trapped. Most of our current knowledge on the assembly of proteins comes from their crystal structures^56^, after taking into account crystal contacts that are not considered. However, crystallography has limitations in determining quaternary structure^57^, as crystallization conditions optimize for perfect order, which is achieved at high protein concentrations and solution additives (salt and crowders). These may push the proteins to form an ordered, homogeneous lattice. In recent years, new methods have been developed that can directly address the assembly state of proteins. Here, we compared 17 different proteins, using four different methods to obtain a more complete picture of their assembly.

A graphical summary of all the data is given in Fig. 6 with detailed descriptions in Table S2. The most visible conclusion from the figure is that it would be wrong to assign a single oligomeric state to proteins. Most proteins appear in more than one state. Moreover, of the selected 17 proteins, none is solely in a monomeric state at all protein concentrations. Second, the predicted multimeric states, as defined in UniProt or the PDB do not consider the complexity of the oligomeric state of the different proteins in solution, with large differences between UniProt predictions and what we found in solution seen for NadK and Can. In other cases, the predictions cover only part of the complex oligomeric sub-states. Conversely, AF Multimer performed better in defining the different multimeric states than the structure-based methods (UniProt, PDB, PiQSi). While AF Multimer does not provide information on the dominant multimeric form (which will very much depend on the solution conditions and protein concentrations), it accurately calculated the potential multimeric states of a protein.

**Figure 6:**
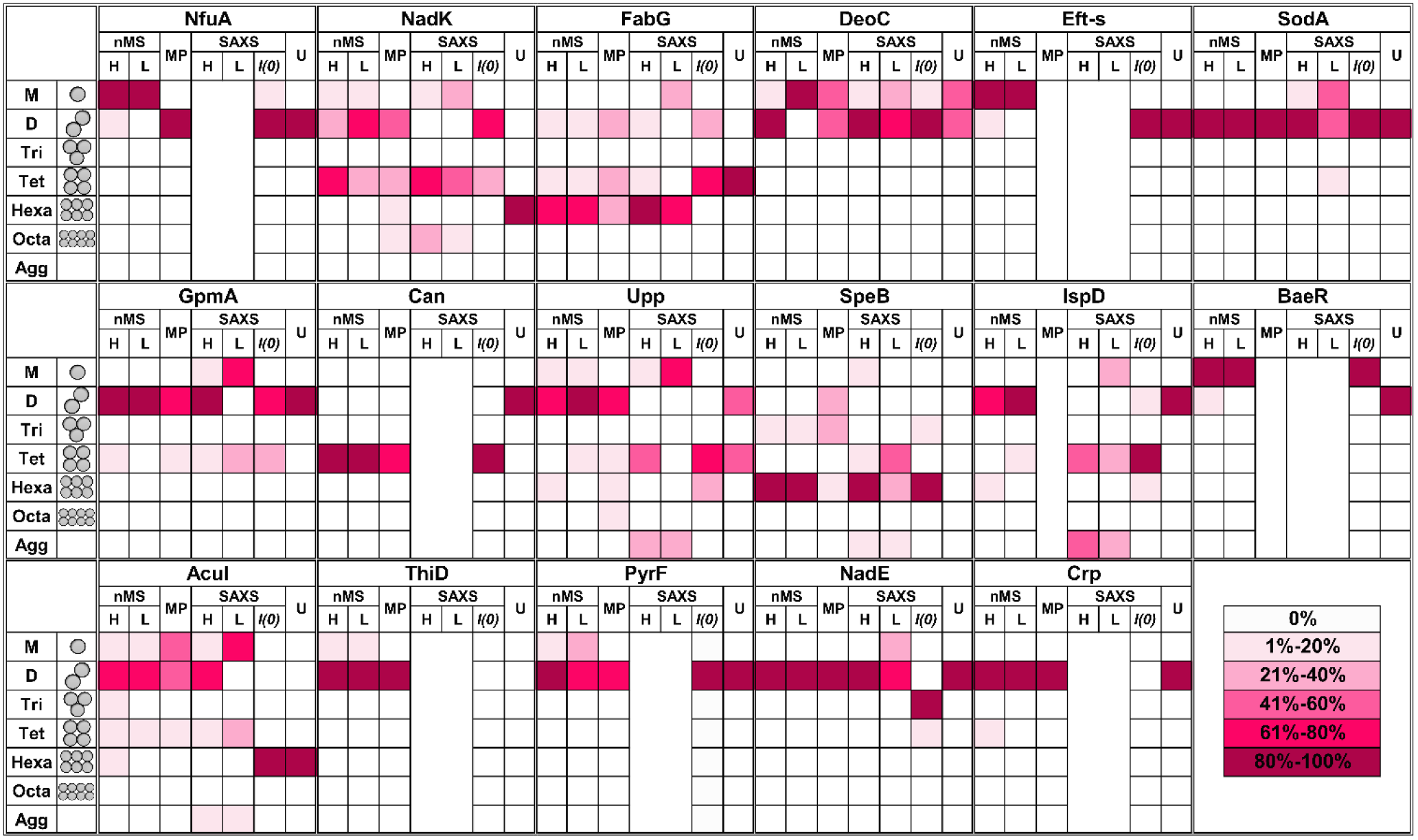
Summary figure for native MS, MP and SAXS results for 17 proteins analyzed in this study. Each box represents percentage of the oligomer in the specific form. The color of the box indicates the percentage, from white – 0% to magenta-80%-100% as presented in the legend on the right panel. Native MS at high (H) concentration was for 40 μM protein in all cases. The low (L) concentration depends on the protein. All MP measurements were done at low concentration (nM). SAXS analysis is shown only for the proteins where the raw data were fitted using OLIGOMER. U = UniProt quaternary structure. For concentrations and SAXS data see Table S1 and Table S2 supplemental SAXS data.

As clearly seen in Fig. 6, the oligomeric state varies even between the different methods used. To rationalize this, we summarize the strong and weak points of each method. SEC elution time is dictated not only by the mass, but also by the shape of the protein. Connecting SEC to a MALS detector, which provides the mass of the out coming protein-peak (Fig. S7). Indeed, Eft-S MALS-SEC determined a MM of 28 kDa as the main peak and 57 kDa as the minor peak (Fig. S7B), in line with the native MS results for this protein (Table S2). Still, MALS-SEC cannot correct for the case where the protein-peak contains a mixture of multiple species, which do not separate in the SEC run. For DeoC (Fig. S7A), MALS-SEC measured a single MM of 47 kDa, while the protein is in monomer (27.5 kDa)/dimer (55 kDa) equilibrium (Fig. 2). Another shortcoming of SEC (or SEC-MALS) is the unknown concentration of the protein during the run, with the protein being diluted as it is proceeding along the column.

Native MS retains non-covalent interactions during electrospray ionization, allowing for the detection of protein and protein complexes with high mass accuracy^58–61^. The samples have to be buffer/electrolyte exchanged prior to native MS analysis as the ionization of the analyte of interest can be suppressed by the non-volatile salts present in the samples (since they tend to outcompete the ionization of the proteins). Also, the non-volatile salts can remain on the proteins after ionization, which can result in different adduct forms of the proteins in addition to the protonated form. MS spectra with more than one adduct form make the data analysis and spectral deconvolution more difficult. The LC-based setup used here enables automated measurements, providing the means to exchange the proteins of interest into MS-compatible conditions just prior to native MS. We chose this method to retain proteins in their initial (MS-incompatible) buffer for as long as possible and limit the time those proteins spend in (MS-compatible) aqueous ammonium acetate solution, reducing the risk of possible biases associated with extended storage in the ammonium acetate solution. It should be considered that the buffer-exchange step results in a dilution of the injected sample, affecting the ability to detect samples at initial sub-micromolar/ low nanomolar protein concentration (although direct nanospray is an alternative for those cases). Furthermore, the dilution can affect the multimeric states, when in rapid equilibrium. Still, Table S2 shows high consistency of the oligomeric state throughout the different protein-concentrations, and the ability to observe the oligomerization state at high resolution (down to single percentiles). The overall trend for all proteins is very clear - the higher the concentration, the higher is the oligomeric state. The jumps are usually by simple multiples, monomer/dimer/tetramer (or trimer/hexamer). DeoC provides an example of a perfectly behaving monomer/dimer protein, with a *K*_D_ of 2 μM. NadK goes from monomer to dimer to tetramer. Interestingly, for some of the proteins the multiple species exist in almost fixed percentiles independent of the protein concentration range examined, for example BSA, FabG and IspD. However, for IspD the results should not be taken at their face-value, as non-reducing SDS-Page analysis (Fig. S4) has shown that about 50% of the protein is in an inter-protein disulfide bonded state making an equilibrium assessment impossible. IspD is the only protein, out of the 17 measured that made substantial inter-protein disulfide bridges.

The next method we explored was SAXS, taking advantage of the high-brilliance EMBL P12 beamline at the PETRA III synchrotron source (DESY, Hamburg)^62^. SAXS provides direct estimation of the MM as calculated from the forward scattering intensity, *I(0)*, in addition to equilibrium fitting using the program OLIGOMER^24^. OLIGOMER utilizes the computed scattering from input high-resolution structures to find a best-fit linear combination of these components to the experimental SAXS data. To calculate the MM from *I(0)* the exact concentration of the protein has to be known, as an error in concentration (or existence of some aggregates or impurities) will directly affect the estimated MM and thus the presumed oligomerization state. Additionally, it must be considered that a mixture of multiple states in solution will provide an average MM that may be misinterpreted as the MM of a single species. Indeed, for many of the studied proteins, dividing the MM estimate based on *I(0)* by the known monomeric MM does not result in an integer (for example NadK gives values of 2.4-2.7 and FabG of 3.5-3.7). Conversely, using OLIGOMER provides the equilibrium composition from the best fit to the data. The drawback is that it requires a reliable structural model as input (which were available for the proteins used here). OLIGOMER uses the complete SAXS curve to model the structure. Fig. 1 shows the difference in the quality of the fit for SodA when considering only a dimeric form of the protein, compared to the equilibrium fit that additionally includes a monomer. Clearly, the monomer/dimer equilibrium provides an improved fit to the experimental data, especially for the lower concentration data sets. In addition, OLIGOMER provides a structural description of the equilibrium. Perhaps the main drawback of this method is the high protein-concentrations (>0.1 mg/ml) needed to obtain high quality data, even at the most powerful SAXS beam-lines.

MP can be used with a large variety of solution conditions, with a minute amount of sample. The disadvantages associated with the MP method is its restriction to MM of >40 kDa (which is reduced to >30 kDa in the 2022 version of the instrument), making us blind towards small proteins. Second, MP works in a specific protein concentration, usually in the range of 100 nM. Still, the method provided high quality results by simple measurements. The MM calculated by MP are off by only a few percent (see for example Fig. 3 panels A versus D, for FabG). Species down to 2% of the total mass can be easily detected (see NadK, Table S2, hexamer and octamer).

Comparing the oligomerization states as determined by the different experimental methods shows that overall there is agreement between them, but with quite big differences in the details (Fig. 6). FabG is a case of interest, as it is a tetramer in the x-ray structure. In SEC the main peak corresponds to a tetramer. The SAXS calculated MM from *I(0)* equals 3.5-3.7 monomer subunits per multimer. However, native MS, MP and SAXS modeling using OLIGOMER clearly show the hexamer to be between the dominant species, together with dimer and tetramer. The oligomerization state mixture was fixes at different times of dilution, different pH values or salt concentrations (Fig. 5S). Finally, unfolding FabG in 8M urea and refolded it resulted in the same mixture of oligomeric states, however, with the dimer and tetramer being now the dominant species (Fig. 3E). The MM measured by native MS and SAXS modeling clearly show that we use the same protein for which the structure has been determined. Therefore, the oligomeric state of this protein in solution is probably a mixture of species, which cannot be seen in the x-ray structure. The observation that the different species abundance is not concentration dependent (as also shown for BSA), but changes after urea unfolding and refolding suggests a high transition state barrier between the oligomeric species, which are therefore in a kinetic trap, which does not obey to mass action equilibrium.

In summary, the different methods used here to evaluate the quaternary structure of proteins emphasize that many proteins have several oligomeric forms. An overview of the characteristics of the different methods, are summarized in Figure S8. While for some proteins there is a dominant quaternary structure, for others there is a dynamic equilibrium between multiple species, and yet others are kinetically trapped into multiple oligomeric forms. Therefore, relying on the x-ray structure to determine the oligomeric structure of the protein will often underestimate the real complexity of the protein in solution. In this sense, using AF provided a positive surprise, as it provided an unbiased picture of the potential oligomeric states, however, without providing a judgment of the dominant specie. This is expected, as the dominant species depends on concentration, pH and solution conditions. This work clearly demonstrates that, together with structure deposition, an additional effort should be made to determine the quaternary structure in solution, and that good and accessible methods and tools now exist to do this.

## Material and Methods

### Cloning and Protein production

Each gene was amplified from BL21 (DE3) bacteria using primers designed for RF cloning and cloned into pET-28-14 His-bdSumo^63^ in adjacent to the sumo-tag. An alanine residue was added before the gene to improve sumo protease cleavage. Expression and purification of the proteins were done as described in^44,45^. Efficient cleavage and elution are achieved by a vector expressing a recombinant protein containing a designed His-tag for specific binding and a sumo protease cleavage site fused to the protein of interest. This allows direct cutting and elution from the Ni-NTA beads, without leaving a trace of the linker protein. In addition, this method allows multiple-proteins to be prepared in parallel. After the standard procedure of Ni-NTA purification (Ni-NTA beads, Merck, cat. 70666-4) and Sumo protease cleavage (in-house production, 1:200 sumo protease 1 mg\ml). The proteins were loaded on Hi-trap Q HP (GE Healthcare, cat.17115401) anion exchange column. FabG and SodA showed poor cleavage from the Ni-NTA column so they were eluted from the Ni-NTA using 300 mM Imidazole and their buffer was exchanged to lower salt concentration (25 mM Tris pH=8). Cleavage was performed for 48-72 hours, which-after the proteins were re-loaded on a Ni-NTA column, which removes the His-tag fused sumo tag that binds the column, while the protein is in the flow-through. Purification was evaluated by SDS-PAGE analysis (ExpressPlus PAGE Gel,15 wells, 4-20%, GeneScript Cat# M42015) with and without β-mercaptoethanol (Genescript, cat: MB01015) added to 10 μg of each protein (Fig. S4). This comes to evaluate inter-disulfide bridges, which would affect the determined oligomeric state. The samples were heated and loaded on gel. Gel was colored by Instant Blue Coomassie Protein Stain (Abcam, ab119211) over-night and then pictured. All proteins have a high degree of purity, with only IspD having a substantial population of inter-disulfide bridged protein. After purification, dialysis against 50 mM HEPES, 50 mM NaCl pH=7.4 was done twice for storage buffer. All samples were snap frozen in liquid nitrogen and stored at −80 °C until further analysis. For BSA analysis Albumine Bovine fraction V (Cat# 1600069, MP Biomedicals, LLC) was used. 50 mg powder was suspended in 1 ml of PBS pH 7.4, after suspension it was dialyzed against 50 mM Hepes 50 mM NaCl pH 7.4 overnight, protein’s concentration was measured in the Nano-drop using 43.82 M^−1^ cm^−1^ as extinction coefficient and 66 kDa as molecular mass of the protein. This procedure was done in all methods that measured BSA.

### Native mass spectrometry

FabG was diluted in 25 mM Tricine, 50 mM NaCl pH 8.5, and all other proteins were diluted in 50 mM HEPES, 50 mM NaCl pH 7.4. After overnight incubation at 4 °C (for all but BSA), the dilutions were measured by online buffer exchange mass spectrometry (OBE-MS) using a Vanquish UHPLC coupled to a Q Exactive Ultra-High Mass Range (UHMR) mass spectrometer (Thermo Fisher Scientific). 1 μL protein was injected onto either a self-packed buffer exchange column (P6 polyacrylamide gel, Bio-Rad Laboratories) or a prototype desalting column from Thermo Fisher Scientific and online buffer exchanged to 200 mM ammonium acetate, pH 6.8 at a flow rate of 100 μL/min^38^. Eluting proteins were ionized via a heated electrospray ionization (HESI) source using a 3.5 kV spray voltage. Mass spectra were recorded over the m/z range 1000 – 14000, at 17500 resolution as defined at 400 m/z. The injection time was set to 200 ms. Voltages applied to the transfer optics were optimized to allow for ion transmission while minimizing unintentional ion activation, with −5 V in-source trapping and a higher-energy collisional dissociation (HCD) of 5 V applied. UniDec software was used for spectral deconvolution and comparison of relative abundances of the oligomeric state^64^.

### Mass Photometry

Microscope coverslips (No. 1.5, 24 × 50 cat# 0107222, Marienfeld) were cleaned by sequential sonication in 50% isopropanol (HPLC grade)/Milli-Q H2O, and Milli-Q H_2_O (5 min each), followed by drying with a clean nitrogen stream. Four gaskets (Reusable culturewell™ gaskets 3mm diam x 1mm depth, cat GBL103250-10EA, Sigma-Aldrich) were cut to 2×2 array, cleaned similarly to the coverslips, and put on top of the coverslip, each sample measured in one well. Immediately prior to mass photometry measurements, protein stocks were diluted in PBS pH 7.4. To focus, fresh buffer was first introduced into the well, and the focal position was identified and secured in place with an autofocus system based on total internal reflection for the entire measurement. For each acquisition, 5 μL of diluted protein (nanomolar concentrations) was added into the well and, following autofocus stabilization, movies of 120 s duration were recorded. Each sample was measured at least three times independently (n ≥ 3). Calibration of the contrast-to-mass conversion was done similarly to the description above, at the same measurement buffer, with the protein Urease (Sigma cat. U7752-1VL), whose oligomer masses are known. All data were acquired using an OneMP mass photometer (Refeyn Ltd, Oxford, UK). Data acquisition was performed using AcquireMP (Refeyn Ltd, v2.2) and data analysis was performed using DiscoverMP (Refeyn Ltd, v2.3.0). Gaussian fit was done using KaleidaGraph software v 4.1 for the AcuI and DeoC proteins due to overlapping areas. For measurements in different salt conditions and varying pH, the calibrations of Urease was done in the same buffer composition of the measurements. For salt conditions: 50mM HEPES pH 7.4, 50 mM HEPES, 500 mM NaCl pH 7.4, 50m M HEPES 1 M NaCl pH 7.4. All buffers were filtered using syringe filters of 0.2 μm (Millipore cat# SLGP033R) before the measurements. For the refolding experiment, 40 μl of 8 M Urea was added to 2 μl FabG, the protein was incubated in this buffer for 30 minutes at room temperature. After the incubation, buffer exchange against PBS was done by using mini GeBAflex tubes (cat#:DO70-6-10, Tivan Biotech). The measurements were done as described above.

### Small Angle X-ray Scattering

Synchrotron radiation X-ray scattering data were collected for all protein samples on the EMBL P12 beamline of the storage ring PETRA III (DESY, Hamburg) using a PILATUS 6M pixel detector (DECTRIS, Switzerland) ^62^. The experimental details of the instruments and derived parameters are listed in Table S1. Forty μl sample were exposed to X-rays while flowing through a temperature-controlled quartz capillary (1.2 mm ID) at 20°C. Forty image frames of 0.045 s exposure time were collected and data from the detector was normalized to the transmitted beam intensity, averaged, buffer subtracted, and placed on an absolute scale relative to water using the SASFLOW pipeline^65^. All data manipulations were performed using PRIMUS*qt* and the ATSAS software package^66^. Where necessary, additional scaling of buffer data sets to minimize mismatch with sample scattering was conducted prior to the subtraction procedure. The forward scattering *I(0)* and radius of gyration, *R*_*g*_ were determined from Guinier analysis^23^, assuming that at very small angles (s ≤ 1.3/*R*_*g*_) the intensity is represented as *I(s)*=*I(0)*exp(-(*sR*_*g*_)2/3)). These parameters were also estimated from the full scattering curves using the indirect Fourier transform method implemented in the program GNOM^67^, along with the distance distribution function *p(r)* and the maximum particle dimensions *D*_*max*_. Molecular masses (*MMs*) of solutes were estimated from *I(0*) by computation of partial specific volume and the contrast between the protein sequence and the chemical components of the solution using in-house procedures. Computation of theoretical scattering intensities from models and PDB files was performed using the program CRYSOL^25^.

### Structure model building using SAXS data

Analysis of the structures present in the solution for each protein sample was conducted using the non-negative linear least-squares routine of the program OLIGOMER^24^, where the experimental scattering intensity *I*_*exp*_*(s)* from a mixture of *K* different particles/components is:

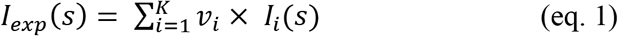

where *v*_*i*_ and *I*_*i*_(*s*) are the volume fraction and the scattering intensity from the *i*-th component. Form-factors were computed from the high-resolution PDB structures available (Table S1), or from homology models from the Swiss-model repository, using FFMAKER^68^. Arrangements of higher oligomers were derived from symmetry mates defined in the PDB files and guided by the PISA server at EBI (https://www.ebi.ac.uk/msd-srv/prot_int/cgi-bin/piserver)^69^, and possible association/dissociation components extracted (eg. dimers and monomers) and form-factors computed. For the generation of a hexameric FabG model the program SASREF^70^ was used, using the monomeric subnunit extracted from the PDB structure (PDB id. 1I01) with P32 symmetry enforced. The form-factors of potential species present in solution were used as input for OLIGOMER and the volume fractions of each component determined through the fitting routine to minimize the discrepancy between the experimental and calculated SAXS curves according to:

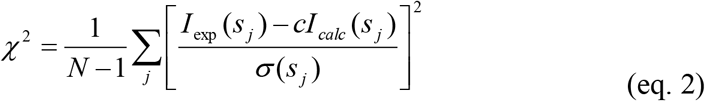

where *N* is the number of experimental points, *c* is a scaling factor and *I*_*calc*_*(s*_*j*_) and *s(s*_*j*_) are the calculated intensity and the experimental error at the momentum transfer *s*_*j*_, respectively. SAXS data have been deposited at the SASBDB (www.sasbdb.org) with accession codes: SASDLR4, SASDLQ4, and SASDLP4.

### Size Exclusion Chromatography

30 μg of each protein was loaded onto a Superdex 200 Increase 10\300 GL column (GE, cat 28-990944) by an Alias™ auto-sampler. The column was pre-equilibrated with PBS pH 7.4 and the proteins were diluted in the same buffer. Proteins for the standard curve were also loaded in the same manner. Standard curve fit (Fig. S1) for known proteins was generated with the following elution volumes (EV) and molecular weights (MM): (EV, MM)- BSA dimer (11.8, 132), BSA monomer (13.51, 66), IFNα2+IFNAR2 (14.5, 43), TEM & BLIP (15.05, 46.7), IFNAR2 (15.6, 24.6), TEM (16.03, 28.9), BLIP (16.7, 17.8), UnaG (16.7, 15.6). The relation between elution volume and the known MM was fitted using an exponential equation.

### SEC-MALS

2-2.5 mg/ml of DeoC or E-fts were loaded into a 100 μl loop on a Superdex 200 Increase 10\300 GL column (GE, cat 28-990944) using the following configuration: ÄKTA Pure 25M, multiple-angle light scattering (MALS) by Wyatt Technology model: DAWN HELEOS II and Optilab TrEX. Mass calibration was done by using 2 mg/ml BSA standard (Bio-Rad, cat.5000206). PBS pH 7.4 was used as isocratic buffer. Buffer and samples were filtered through 0.1um filter system.

### AlphaFold analysis

AlphaFold2^37,55^ was implemented by locally running an adapted code written by ColabFold^71^. All runs used the five model parameters presented in CASP14, with no templates or Amber relaxation and performing three recycles. Multiple-sequence alignments were generated through the MMseqs2 API server^72–74^. For Rosetta relaxation, scoring was performed using the ref15 energy function^75^. The relaxation comprises four iterations of sidechain packing and harmonically constrained whole-protein minimization on the input structure. XML is provided in Supplementary Table S3.

#### Supporting Information

Supplementary data of SAXS measurements – Table S1 and for all proteins measurements – Table S2 as well as supplementary figures.

#### Funding Sources

This research was supported by the Israel Science Foundation grant No. 1268/18 (GS). The SAXS experiment at the P12 EMBL beamline at the PETRA III storage ring of DESY synchrotron, Hamburg, Germany, was supported by iNEXT project number 653706, funded by the Horizon 2020 Program of the European Union. Native mass spectrometry measurements were provided by the NIH-funded Resource for Native Mass Spectrometry Guided Structural Biology at The Ohio State University (NIH P41 GM128577, (VW)).

## Notes

**Competing Interest Statement:** none.

### Competing Interest Statement

The authors have declared no competing interest.

