## Supplementary Figures for "Protein Quaternary Structures in Solution are a Mixture of Multiple forms"

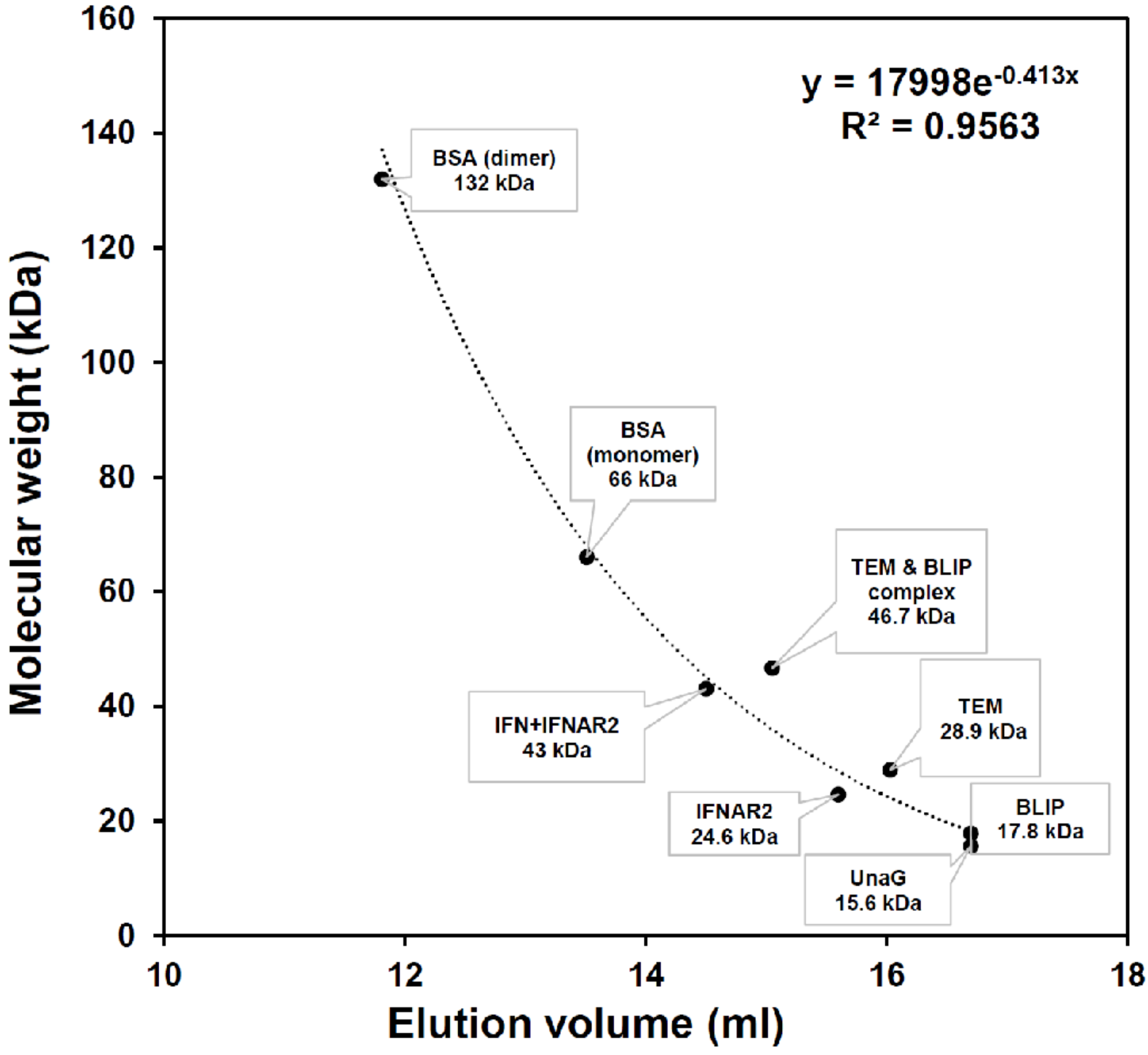


**Figure S1- Standards calibration curve fit for known proteins elution volumes and molecular weights.** The proteins (from largest to smallest- elution volume (ml), Mw (kDa)). BSA dimer (11.8, 132), BSA monomer (13.51, 66), IFN+IFNAR2 (14.5, 43), TEM & BLIP (15.05, 46.7), IFNAR2 (15.6, 24.6), TEM (16.03, 28.9), BLIP (16.7, 17.8), UnaG (16.7, 15.6). The data fitted best an exponential, which was used to calculate the MW of unknown proteins.

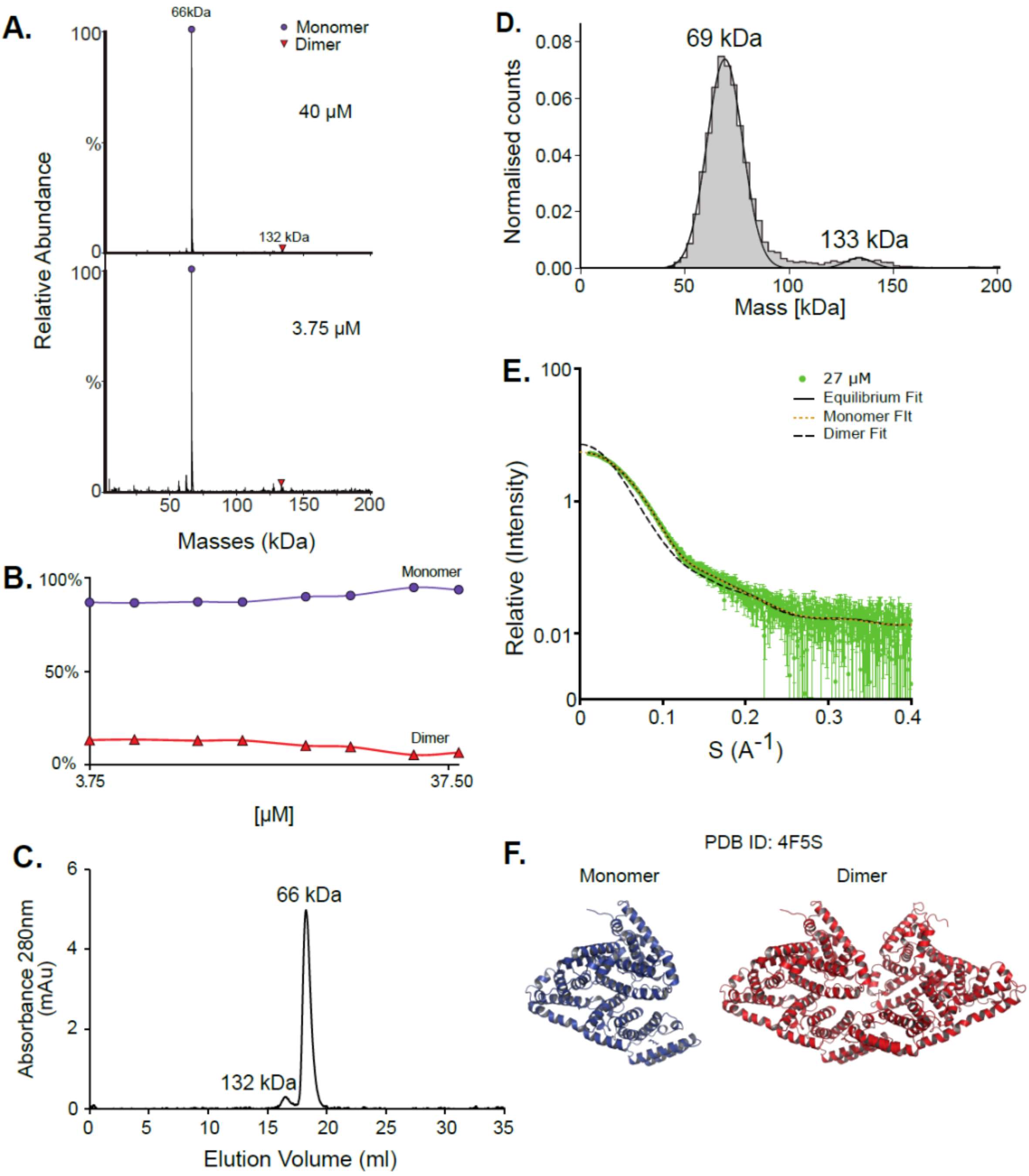


**Figure S2- BSA is mostly a monomer with some small dimeric fraction.** Measurements of BSA, a well known protein, in all the different methods results in similar quaternary structure- mostly a monomer with small dimeric fraction. **A.** Native MS results shows one main peak that corresponds to a 66 kDa monomer and a small peak of 132 kDa dimer in a ration that is not concentration dependent. **B.** Native MS in a range of protein concentrations, 3.75 µM - 40 µM, shows the proteins oligomeric state to be independent on the concentration (see also fig. S3). **C**. SEC analysis shows two peaks- the small one eluted at 11.8 ml corresponds to 132 kDa (a dimer) and the second, main one, eluted at 13.5 ml corresponds to the monomeric form of BSA at 66 kDa. **D**. Mass photometry measurements of the protein show masses that fit a monomer and a dimer- 69 kDa and 133 kDa. **E.** SAXS measurements were done in one concentration of 27 µM and shows that more than 90% of the protein is in monomeric form. SAXS equilibrium fitting using the program OLIGOMER and PDB id: 4F5S shows that the data is well fitted with the equilibrium and monomer but poorly with the dimeric fit (black dashed line). **F.** Assemblies of BSA using OLIGOMER and the fit.

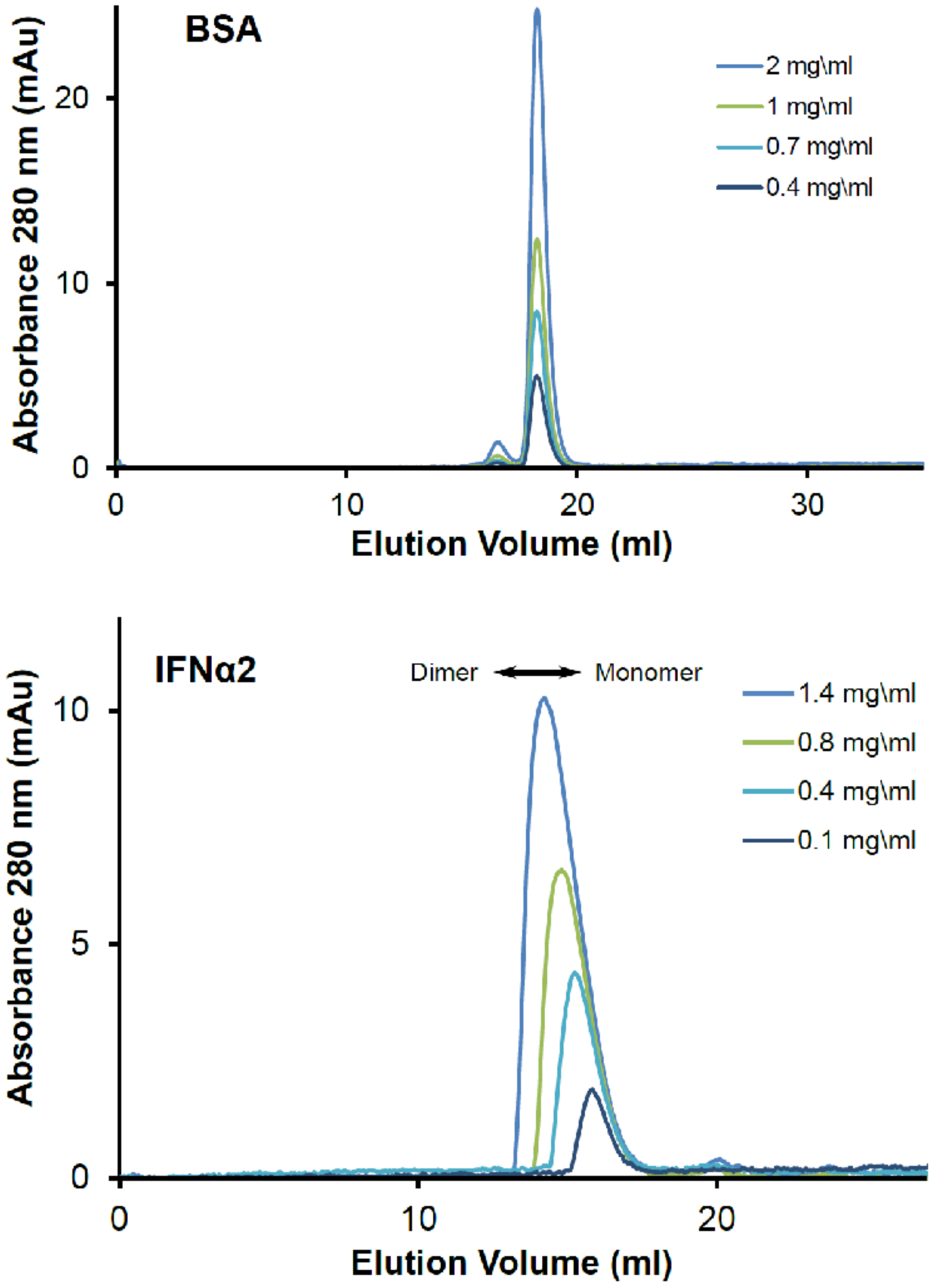


**Figure S3- SEC concentration-dependent elution of BSA and IFNα2.** SEC analysis of BSA at 0.4-2 mg\ml and IFNα2 0.1-1.4 mg\ml shows that BSA's elutes at the same volume, whereas c elution volume decreases with increasing concentration. This suggests a concentration dependent oligomerization of IFNα2.

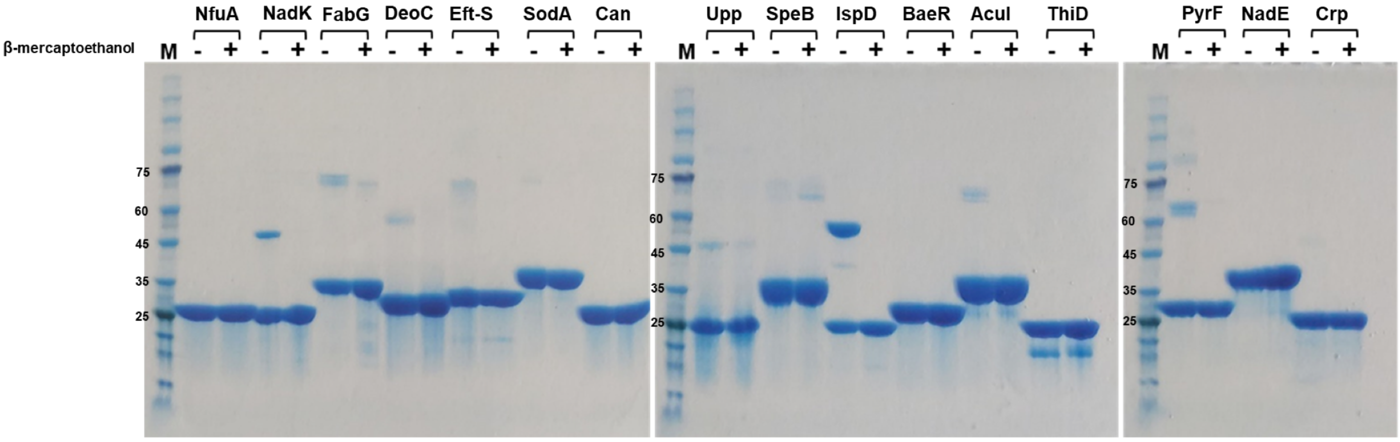


| Protein's name | # of cys residues |
|---|---|
| NfuA | 4 |
| NadK | 6 |
| FabG | 0 |
| DeoC | 4 |
| Eft-s | 2 |
| SodA | 0 |
| Can | 5 |
| Upp | 1 |

| Protein's name | # of cys residues |
|---|---|
| SpeB | 4 |
| IspD | 5 |
| BaeR | 4 |
| AcuI | 3 |
| ThiD | 3 |
| PyrF | 3 |
| NadE | 3 |
| CRP | 3 |

**Figure S4- SDS-PAGE analysis of all proteins with and without reducing agent- β-mercaptoethanol.** The gel represents each protein with and without the addition of β-mercaptethanol prior to the heating and loading to the gel. The table represents the number of cysteine residues in each of the proteins. The gel shows that the only protein where inter-disulfide bridges were formed is IspD, while for the other proteins the dominatnt form is the same with and without reducing.

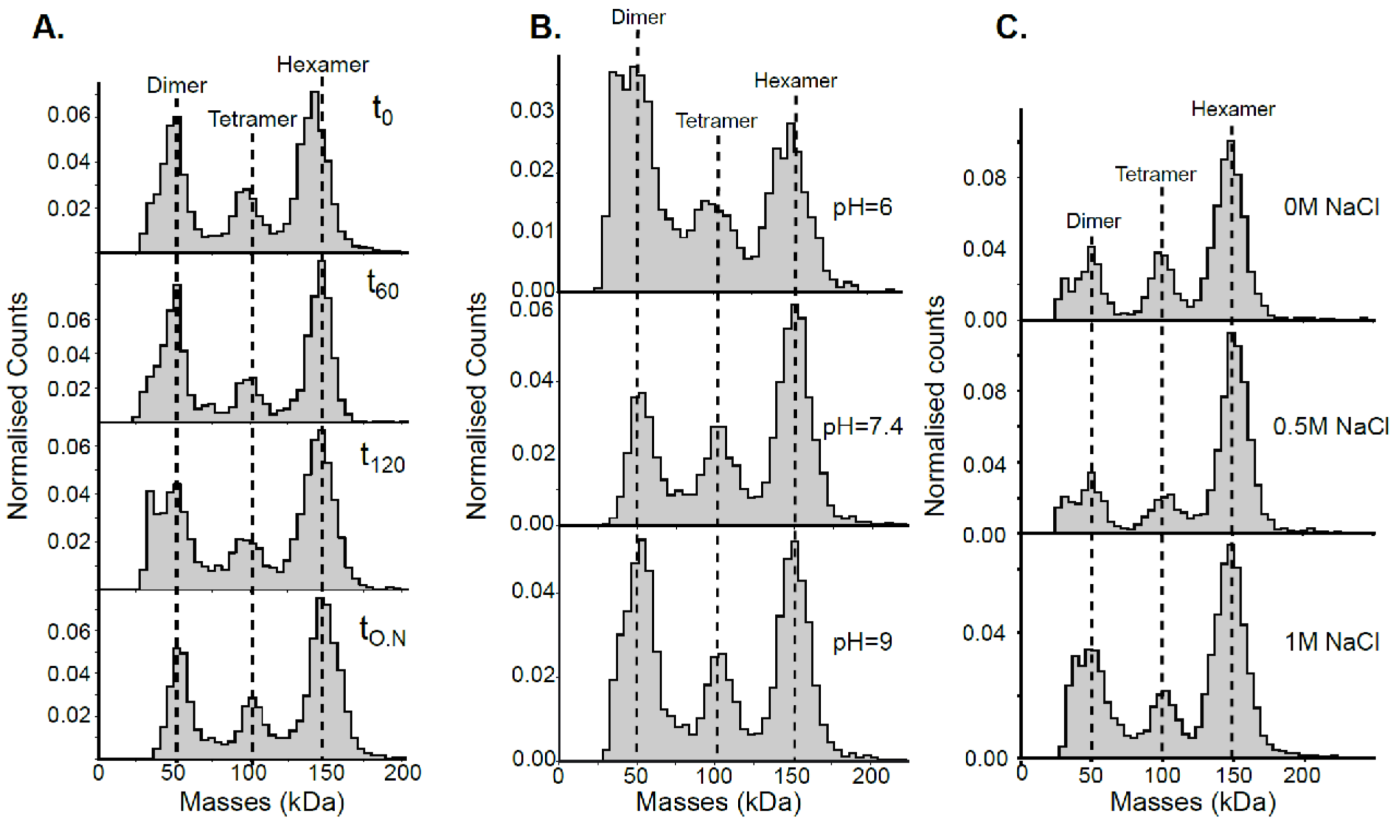


**Figure S5- FabG oligomerizations state equilibrium is not affected by protein-dilution or buffer**. FabG oligomerization state was determined at a concentration of 38 nM by MP. A. Measurements of time points after dilution from 60 µM of FabG shows similar oligomeric states at all times. B. FabG oligomerization states at pH=6 (50mM Sodium Citrate, 50mM NaCl pH=6), pH=7.4 (PBS) and pH=9 (50mM Tricine , 50mM NaCl pH=9). Overall, the changes in the fraction of the different oligomeric states between pH 6-9 are small. C. Salt dependence of the oligomerization state of FabG: 0 M, 500 mM and 1M NaCl in 50mM HEPES buffer, pH 7.4 were used. FabG has shown a similar ration between hexameric, tetrameric and dimeric forms at all three salt concentrations.

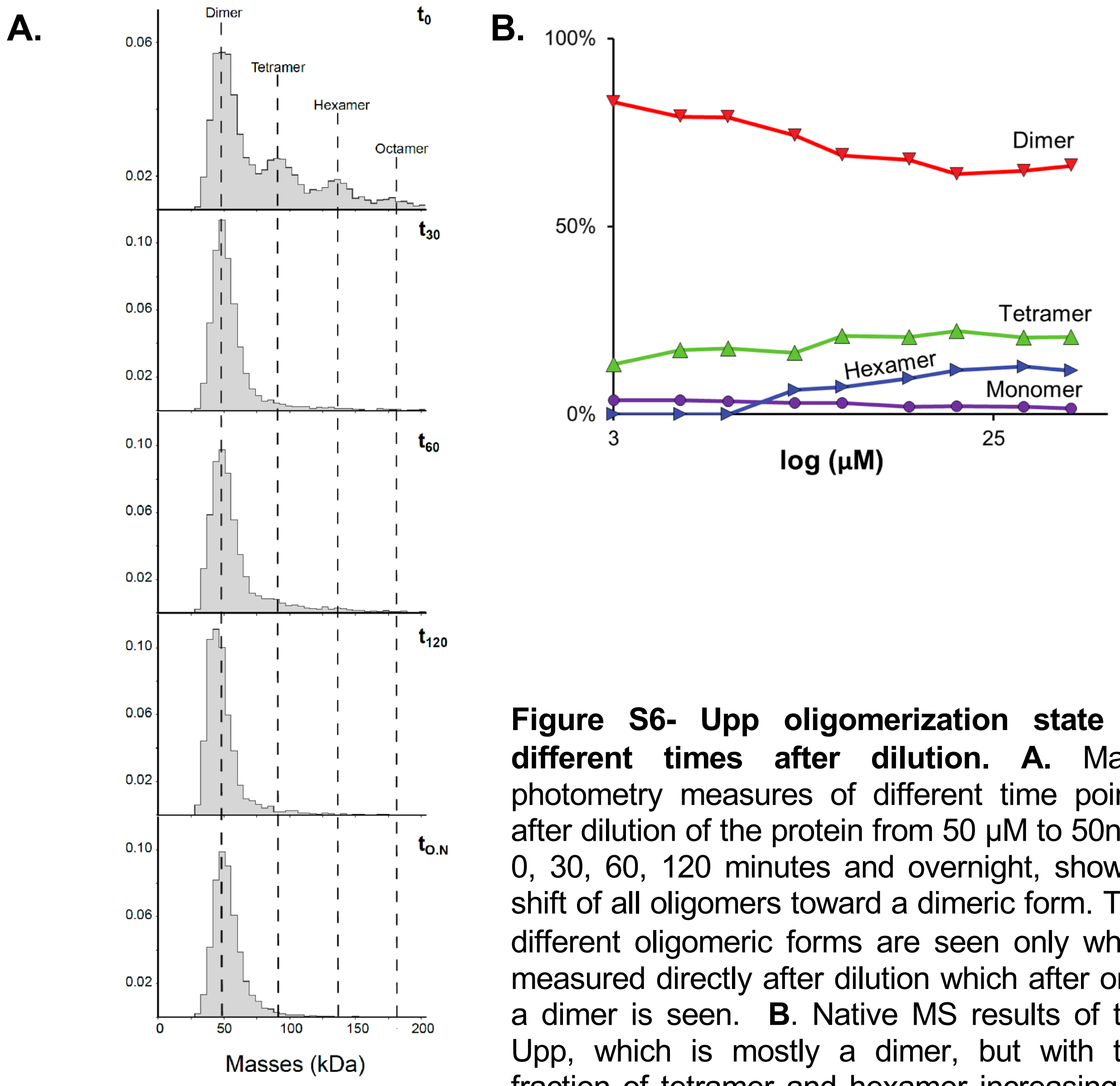


**Figure S6- Upp oligomerization state at different times after dilution. A.** Mass photometry measures of different time points after dilution of the protein from 50 µM to 50nM: 0, 30, 60, 120 minutes and overnight, show a shift of all oligomers toward a dimeric form. The different oligomeric forms are seen only when measured directly after dilution which after only a dimer is seen. **B**. Native MS results of the Upp, which is mostly a dimer, but with the fraction of tetramer and hexamer increasing at higher protein concentrations

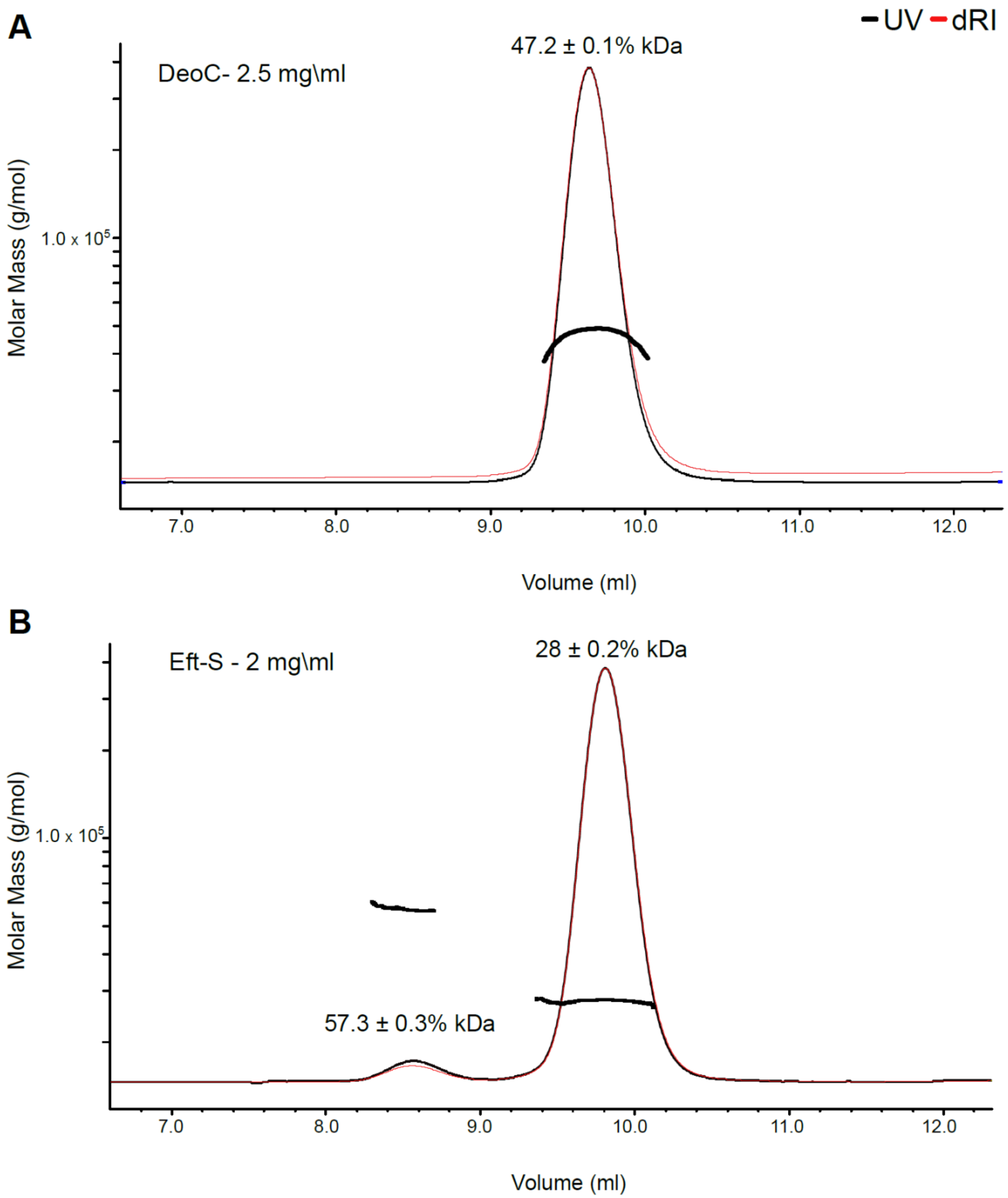


**Figure S7- SEC-MALS of E-fts and DeoC. A.** DeoC is eluted as a single peak, with MALS-detector measureing a MM of 47.2 kDa. As this MM does not corresponds to a monomer (27.7 kDa) or a dimer ( 55 kDa), we conclude that the peak is a mixture of both. **B.** Eft-S is eluted in two peaks, a minor dimeric peak corresponding to 57.3 kDa and a major monomeric peak corresponding to 28 kDa.

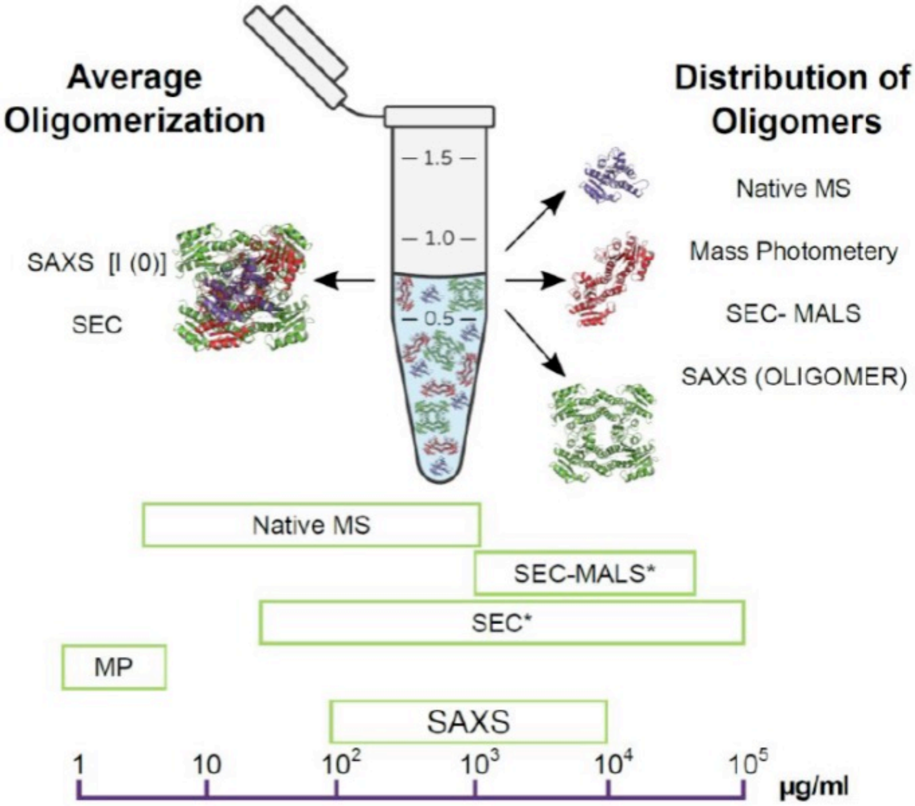


**Figure S8- Graphical summary representation of the different methods to determine oligomerization.** Comparing the different methods for determining oligomerization composition of a protein. Each method is suitable for different protein concentrations. *I(0)* from SAXS as well as SEC give information of the average oligomerization state, whereas, native MS, MP, SEC-MALS (depending on the equilibrium of the different oligomers), and SAXS (by using OLIGOMER) determine the distribution of the oligomers in solution. The (*) in the SEC methods represent the injected concentration that is diluted during the run of the SEC. The ruler of µg/ml represents protein concentrations applicable for the different methods.
