## Supplementary Table S1 for "Protein Quaternary Structures in Solution are a Mixture of Multiple forms"

***Summary table of small-angle X-ray scattering results***

| Sample | *Conc. (mg/ml)* | *R*_g_ (Å) | *d*_max_ (Å) | *M_r_* from *I*(0) (Da) (ratio to predicted value) |
| --- | --- | --- | --- | --- |
| SodA | 0.25 | 21.2 ± 0.2 | 70 ± 5 | 52542 (2.3) |
|  | 0.51 | 22.7 ± 0.1 | 75 ± 5 | 48500 (2.1) |
|  | 1.01 | 22.7 ± 0.1 | 75 ± 5 | 45806 (2.0) |
|  | 2.03 | 22.7 ± 0.1 | 72 ± 5 | 45806 (2.0) |
| DeoC | 0.26 | 25.2 ± 0.3 | 85 ± 5 | 55606 (2.0) |
|  | 0.52 | 25.7 ± 0.1 | 85 ± 5 | 54216 (2.0) |
|  | 1.04 | 26.1 ± 0.1 | 85 ± 5 | 51436 (1.9) |
|  | 2.08 | 26.0 ± 0.1 | 80 ± 5 | 50046 (1.8) |
| FabG | 0.24 | 33.2 ± 0.1 | 100 ± 5 | 89174(3.5) |
|  | 0.48 | 33.7 ± 0.1 | 100 ± 5 | 91961 (3.6) |
|  | 0.95 | 33.9 ± 0.1 | 105 ± 5 | 91961 (3.6) |
|  | 1.90 | 34.0 ± 0.2 | 106 ± 5 | 93354 (3.7) |
| NadK | 0.25 | 36.2 ± 0.3 | 11.5 ± 5 | 77202 (2.4) |
|  | 0.5 | 3.7 ± 0.2 | 12 ± 5 | 81413 (2.5) |
|  | 1.0 | 38.4 ± 0.1 | 12 ± 5 | 87028 (2.7) |
|  | 2.0 | 39.8 ± 0.1 | 12.5 ± 5 | 87828 (2.7) |

**TABLE SAXS1A**

***Small-angle X-ray scattering parameters and results SodA, DeoC, FabG***

| (*a*) Sample details | |  |  |
| --- | --- | --- | --- |
|  | SodA | DeoC | FabG |
| Organism | *Escherichia coli (strain K12)* | *Escherichia coli (strain K12)* | *Escherichia coli (strain K12)* |
| Source | *Escherichia coli BL21 (DE3)* | *Escherichia coli BL21 (DE3)* | *Escherichia coli BL21 (DE3)* |
| UniProt sequence ID (residues in construct) | P00448 | P0A6L0 | P0AEK2 |
| Extinction coefficient ε (280 nm, 0.1% w/v) | 1.893 | 0.523 | 0.452 |
| Partial specific volume 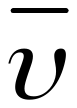 (cm^3^ g^-1^) | 0.737 | 0.741 | 0.742 |
| Mean solute and solvent scattering length densities and mean scattering contrast 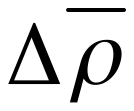 (*ρ_protein_-ρ_solvent_)* (10^10^ cm^-2^) | 2.87 (12.297-9.429) | 2.81 (12.238-9.429) | 2.80 (12.231-9.429) |
| Molecular mass *M* from chemical composition (monomer) (Da) | 22950 | 27619 | 25377 |
| Sample concentration (mg ml^-1^) [A280nm] | 0.25-2.0 | 0.26-2.08 | 0.24-1.90 |
| Sample volume (ul) | 40 | | |
| Solvent composition | 50 mM HEPES pH 7.2 | | |
| (*b*) SAS data collection parameters | |  |  |
| Instrument/Data processing | EMBL P12 (PETRA-III, DESY, Hamburg) with Pilatus6M detector (Blanchet et al. 2015) | | |
| Wavelength (Å) | 1.24 | | |
| Beam geometry (size, sample-to-detector distance) | 0.12 × 0.25 mm^2^, 3.0 m | | |
| *s*-measurement range (Å^-1^) | 0.002-0.5 | | |
| Absolute scaling method | Comparison with scattering from 1.2 mm pure H_2_O | | |
| Basis for normalization to constant counts | To transmitted intensity by beam-stop counter | | |
| Method for monitoring radiation damage | Frame comparison | | |
| Exposure time, number of exposures | 1.8 s (40 × 0.045 s) | | |
| Sample temperature (ºC) | 20 | | |
| (*c*) Software employed for SAS data reduction, analysis and interpretation | |  |  |
| SAS data reduction | *I(s)* versus *s* using *RADAVER* (ATSAS 2.8.3; Petoukhov et al., 2012), solvent subtraction using *PRIMUSqt* (ATSAS 2.8.3; Petoukhov et al., 2012) | | |
| Calculation of ε from sequence | *ProtParam* (Gasteiger et al., 2005) | | |
| Calculation of 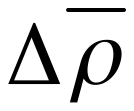 and 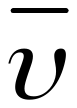 values from chemical composition | Direct Calculation (in-house routines) (Fraser et al. 1978) | | |
| Basic analyses: Guinier, *P*(*r*), scattering particle volume (*V*_P_) | *PRIMUSqt* from ATSAS 2.8.3 (Petoukhov *et al*., 2012) | | |
| Equilibrium analysis | OLIGOMER (Konarev et al., 2003) | | |
| Atomic structure modelling | CRYSOL (Svergun et al., 1995), SASREF (Petoukhov et al., 2005) | | |
| Molecular graphics | PyMOL v2.3 MacOS 10.13.6 | | |
| (*d*) Structural parameters^a^ | |  |  |
| Guinier Analysis | SodA | DeoC | FabG |
| *I*(0) (cm^-1^) | 0.034 ± 0.001 | 0.036 ± 0.001 | 0.067 ± 0.001 |
| *R*_g_ (Å) | 22.8 ± 0.1 | 25.6 ± 0.1 | 35.5 ± 0.1 |
| *q-*range (Å^-1^) | 0.012-0.057 | 0.011-0.051 | 0.016-0.036 |
| *M_r_* from *I*(0) (Da) (ratio to predicted value) | 45806 (2.0) | 50046 (1.8) | 93354 (3.7) |
| *P*(*r*) analysis | SodA | DeoC | FabG |
| *I*(0) (cm^-1^) | 0.034 ± 0.001 | 0.036 ± 0.001 | 0.066 ± 0.001 |
| *R*_g_ (Å) | 22.7 ± 0.1 | 26.0 ± 0.1 | 34.0 ± 0.1 |
| *d*_max_ (Å) | 72.2 ± 5 | 80.0 ± 5 | 106 ± 5 |
| *q-*range (Å^-1^) | 0.012-0.287 | 0.011-0.287 | 0.016-0.287 |
| χ^2^ (total estimate from *GNOM*) | 1.0 (0.94) | 1.1 (0.93) | 1.1 (0.83) |
| *M_r_* from *I*(0) (Da) (ratio to predicted value) | 45967 (2.0) | 50477 (1.8) | 91264 (3.6) |
| Volume(*V*_P_) (Å^3^) | 53629 | 61429 | 212014 |
| *M_r_* from *V*_P_ (Da) (ratio to predicted value) | 33518 (1.5) | 38393 (1.4) | 132509 (5.2) |
| (*e*) Equilibrium modeling results | |  |  |
| *OLIGOMER* fitting | SodA | DeoC | FabG |
| Starting crystal structures | 1D5N | 1KTN | 1I01 |
| Multimers used | Tetramer, dimer, monomer | Dimer, monomer | Hexamer, tetramer, dimer, monomer |
| *q-*range for fitting (Å) | 0.014-0.359 | 0.014-0.359 | 0.014-0.359 |
| χ^2^, *CORMAP* *P* value | 1.1-1.2 (0.005-0.072) | 1.2-1.7 (0.000) | 1.2-1.5 (0.000-0.260) |
| (f) SASBDB IDs for data and models |  |  |  |
|  | SodA | DeoC | FabG |
|  | SASDLR4 | SASDLQ4 | SASDLP4 |

^a^parameters reported for highest sample concentration

**TABLE SAXS1B**

***Small-angle X-ray scattering parameters and results for NadK***

| (*a*) Sample details | |
| --- | --- |
|  | NadK |
| Organism | *Escherichia coli (strain K12)* |
| Source | *Escherichia coli BL21 (DE3)* |
| UniProt sequence ID (residues in construct) | P0A7B3 |
| Extinction coefficient ε (280 nm, 0.1% w/v) | 0.750 |
| Partial specific volume 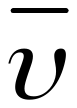 (cm^3^ g^-1^) | 0.742 |
| Mean solute and solvent scattering length densities and mean scattering contrast 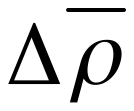 (*ρ_protein_-ρ_solvent_)* (10^10^ cm^-2^) | 2.79 (12.220-9.429) |
| Molecular mass *M* from chemical composition (monomer) (Da) | 32566 |
| Sample concentration (mg ml^-1^) [A280nm] | 0.25-2.0 |
| Sample volume (ul) | 40 |
| Solvent composition | 50 mM HEPES pH 7.2 |
| (*b*) SAS data collection parameters | |
| Instrument/Data processing | EMBL P12 (PETRA-III, DESY, Hamburg) with Pilatus6M detector (Blanchet et al. 2015) |
| Wavelength (Å) | 1.24 |
| Beam geometry (size, sample-to-detector distance) | 0.12 × 0.25 mm^2^, 3.0 m |
| *s*-measurement range (Å^-1^) | 0.002-0.5 |
| Absolute scaling method | Comparison with scattering from 1.2 mm pure H_2_O |
| Basis for normalization to constant counts | To transmitted intensity by beam-stop counter |
| Method for monitoring radiation damage | Frame comparison |
| Exposure time, number of exposures | 1.8 s (40 × 0.045 s) |
| Sample temperature (ºC) | 20 |
| (*c*) Software employed for SAS data reduction, analysis and interpretation | |
| SAS data reduction | *I(s)* versus *s* using *RADAVER* (ATSAS 2.8.3; Petoukhov et al., 2012), solvent subtraction using *PRIMUSqt* (ATSAS 2.8.3; Petoukhov et al., 2012) |
| Calculation of ε from sequence | *ProtParam* (Gasteiger et al., 2005) |
| Calculation of 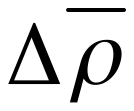 and 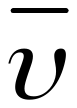 values from chemical composition | Direct Calculation (in-house routines) (Fraser et al. 1978) |
| Basic analyses: Guinier, *P*(*r*), scattering particle volume (*V*_P_) | *PRIMUSqt* from ATSAS 2.8.3 (Petoukhov *et al*., 2012) |
| Equilibrium analysis | OLIGOMER (Konarev et al., 2003) |
| Atomic structure modelling | CRYSOL (Svergun et al., 1995), SASREF (Petoukhov et al., 2005) |
| Molecular graphics | PyMOL v2.3 MacOS 10.13.6 |
| (*d*) Structural parameters^a^ | |
| Guinier Analysis | NadK |
| *I*(0) (cm^-1^) | 0.0630 ± 0.001 |
| *R*_g_ (Å) | 40.3 ± 0.2 |
| *q-*range (Å^-1^) | 0.1474-0.3143 |
| *M_r_* from *I*(0) (Da) (ratio to predicted value) | 88432 (2.7) |
| *P*(*r*) analysis | NadK |
| *I*(0) (cm^-1^) | 0.0626 ± 0.001 |
| *R*_g_ (Å) | 39.8 ± 0.1 |
| *d*_max_ (Å) | 12.5 ± 5 |
| *q-*range (Å^-1^) | 0.1474-2.8732 |
| χ^2^ (total estimate from *GNOM*) | 1.2 (0.89) |
| *M_r_* from *I*(0) (Da) (ratio to predicted value) | 87828 (2.7) |
| Volume(*V*_P_) (Å^3^) | 259247 |
| *M_r_* from *V*_P_ (Da) (ratio to predicted value) | 162029 (5.0) |
| (*e*) Equilibrium modeling results | |
| *OLIGOMER* fitting | NadK |
| Starting crystal structures | 4HAO |
| Multimers used | 8-mer, tetramer, dimer, monomer |
| *q-*range for fitting (Å) | 0.014-0.359 |
| χ^2^, *CORMAP* *P* value | 1.2-1.5 (0.00-0.01) |
| (f) SASBDB IDs for data and models |  |
|  | NadK |
|  | SASDMT3 |

^a^parameters reported for highest sample concentration
