## Supplementary Table S2 for "Protein Quaternary Structures in Solution are a Mixture of Multiple forms"

#### Table S2 - Results summary table of 17 *E.Coli* proteins

##### NfuA- P63020

| Method | MS conc. | MS, kDa | MP, kDa (%) | SAXS, kDa | SEC, kDa | Swiss model calc. | Uniprot (kDa) | PDB | Protein abundance (copy | |
|---|---|---|---|---|---|---|---|---|---|---|
| Concentrations | 0.4 - 40 µM | | 21 nM | 9-72 µM<br>0.2-1.5mg\ml | 14.33 µM | | | NA | Ref.1 | Ref.2 |
| Monomer | | 20.937 | below treshold | *I(0)* = 39.2 (σ=3.5) | 31 (24-40) | Monomer | 20.930 | | 10200 | 3850 |
| Dimer | | | 37 (95%) | | | | Homodimer | | | |
| Graph | 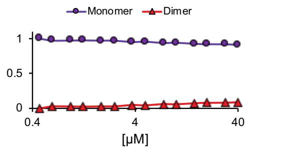 | | 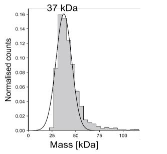 | | 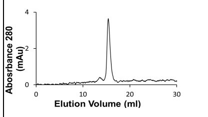 | | | | highly expressed protein | |



##### NadK- P0A7B3

| Method | MS conc. | MS, kDa | MP, kDa (%) | SAXS, kDa | SEC, kDa | Swiss model calc. | Uniprot (kDa) and oligomerization | PDB | Protein abundance (copy number per cell) | |
|---|---|---|---|---|---|---|---|---|---|---|
| Concentrations | 1.25 - 40 µM | | 88 nM | 8-61 µM<br>0.25-2mg\ml | 9.2 µM | | | 4HAO (similar in 82.5%) | Ref.1 | Ref.2 |
| Monomer | | 32.51 | | | | | 32.57 | | NA | |
| Dimer | | 65.01 | 63 (46%) | *I(0)* =83.5 (σ=5.2) | | Dimer | | | | |
| Tetramer | | 130.02 | 123 (39%) | | 88 (69-113) | | | | | |
| Hexamer | | | 190 (2%) | | | | Homohexamer | | | |
| Octamer | | | 248 (2%) | | | | | | | |
| Graph | 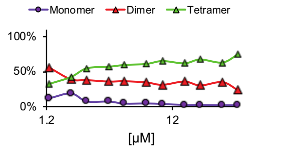 | | 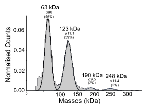 | OLIGOMER software fit: 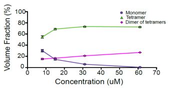 | 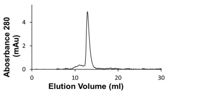 | | | | | |



##### FabG- P0AEK2

| Method | MS conc. | MS, kDa | MP, kDa (%) | SAXS, kDa | SEC, kDa | Swiss model calc. | Uniprot (kDa) and oligomerization | PDB | Protein abundance (copy number per cell) | |
|---|---|---|---|---|---|---|---|---|---|---|
| concentrations | 1.87 - 40 µM | | 38 nM | 9-74 µM<br>0.24-1.9 mg\ml | 11.74 µM | | | 1I01 - homotetramer 1q7b- homotetramer 1q7c- homotetramer | Ref.1 | Ref.2 |
| Monomer | | | | | | | 25.56 | | 13800 | 6053 |
| Dimer | | 50.89 | 51 (32%) | | | | | | highly expressed protein | |
| Tetramer | | 102 | 100 (22%) | *I(0)* =91.6 (σ=1.8) | 101 (79-129) | Homotetramer | Homotetramer | | | |
| Hexamer | | 152.68 | 148 (38%) | | | | | | | |
| Graph | 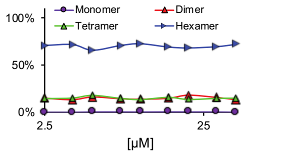 | | 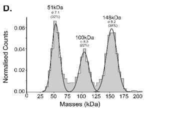 | OLIGOMER software fit: 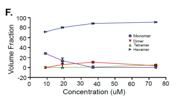 | 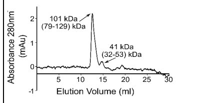 | | | | | |



##### DeoC - P0A6L0

| Method | MS conc. | MS, kDa | MP, kDa (%) | SAXS, kDa | SEC, kDa | Swiss model calc. | Uniprot (kDa) and oligomerization | PDB | Protein abundance (copy number per cell) | |
|---|---|---|---|---|---|---|---|---|---|---|
| concentrations | 0.17 - 40 µM | | 53 nM | 9-75 µM<br>0.3-2 mg\ml | 10.8 µM | | | 1KTN 1JCJ 5EMU | Ref.1 | Ref.2 |
| Monomer | | 27.69 | 36 (68%) fitted: 35(52%) | | 45 (35-57) | Monomer and homodimer | 27.74<br>Monomer and | | 67100 | 6908 |
| Dimer | | 55.38 | 52 (58%) fitted: 52(48%) | *I(0)* =52.8 (σ=2.5) | | | | | highly expressed protein | |
| Graph | 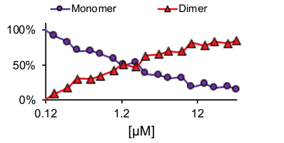 | | 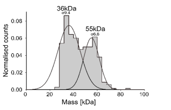 | OLIGOMER software fit: 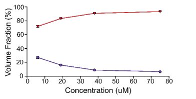 | 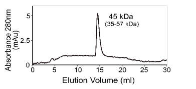 | | | | | |

### Eft-S - P0A6P1

| Method | MS conc. | MS, kDa | MP, kDa (%) | SAXS, kDa | SEC, kDa | Swiss model calc. | Uniprot (kDa) and oligomerization | PDB | Protein abundance (copy number per cell) | |
|---|---|---|---|---|---|---|---|---|---|---|
| concentrations | 0.94 - 40 μM | | | 7-56uM<br>0.2-1.7 mg\ml | 9.9 μM | | | | Ref.1 | Ref.2 |
| Monomer | | 30.36 | | $I(0)$ = 64.7 (σ=5.2) | 41 (32-53) | monomer | 30.29 | | 66000 | 14933 |
| Dimer | | | | | | Dimer- hetromer | Dimer | | | |
| Tetramer | | | | | | | | | | |
| Graph | 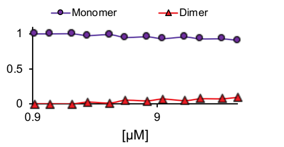 | | NA<br>Under the detection limit of MP | | 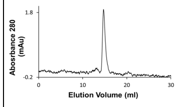 | | | 1EFU | highly expressed protein | |

### SodA- P00448

| Method | MS conc. | MS, kDa | MP, kDa (%) | SAXS, kDa | SEC, kDa | Swiss model calc. | Uniprot (kDa) and oligomerization | PDB | Protein abundance (copy number per cell) | |
|---|---|---|---|---|---|---|---|---|---|---|
| concentrations | 0.94 - 40 μM | | 150 nM | 11-88 μM<br>0.25-2 mg\ml | 1.36 μM | | | | Ref.1 | Ref.2 |
| Monomer | | | | | 36 (28-45) | | 22.97 | | 36900 | 11930 |
| Dimer | | 46.04 | 52 (97%) | $I(0)$ = 48.2 (σ=3.2) | | Dimer | Homodimer | | | |
| Graph | 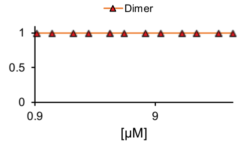 | | 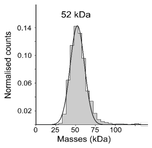 | OLIGOMER software fit: 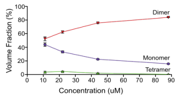 | 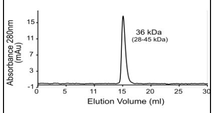 | | | 1IXB 1D5N 1VEW | highly expressed protein | |

### GpmA- P62707

| Method | MS conc. | MS, kDa | MP, kDa (%) | SAXS, kDa | SEC, kDa | Swiss model calc. | Uniprot (kDa) and oligomerization | PDB | Protein abundance (copy number per cell) | |
|---|---|---|---|---|---|---|---|---|---|---|
| concentrations | 0.156 - 40 μM | | 128 nM | 10-77 μM<br>0.3-2 mg\ml | 10.55 μM | | | | Ref.1 | Ref.2 |
| Monomer | | | | | | | 28.43 | | 14400 | 6169 |
| Dimer | | 56.99 | 59 (80%) | $I(0)$ = 68.7 (σ=4.7) | | Homodimer | Homodimer | | | |
| Tetramer | | 114.00 | 116 (4%) | | | | | | | |
| Graph | 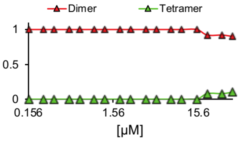 | | 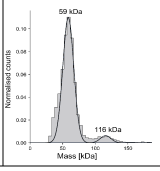 | | 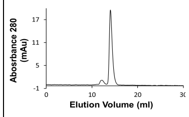 | | | 1E58 | highly expressed protein | |

### Can- P61517

| Method | MS conc. | MS, kDa | MP, kDa (%) | SAXS, kDa | SEC, kDa | Swiss model calc. | Uniprot (kDa) and oligomerization | PDB | Protein abundance (copy number per cell) | |
|---|---|---|---|---|---|---|---|---|---|---|
| concentrations | 0.938 - 40 μM | | 60 nM | 10-83 μM<br>0.3-2 mg\ml | 12 μM | | | | Ref.1 | Ref.2 |
| Monomer | | 25.10 | | | | | 25.1 | | 4940 | 1611 |
| Dimer | | | | $I(0)$ = 86.4 (σ=5.3) | | | Homodimer | | | |
| Trimer | | | | | 77 (60-98) | | | | | |
| Tetramer | | 100.40 | 99 (75%) | | | Homotetramer | | | | |
| Graph | 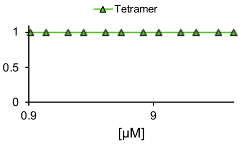 | | 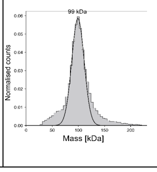 | | 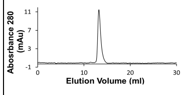 | | | 4ZNZ 1T75 | highly expressed protein | |

#### Upp- P0A8F0

| Method | MS conc. | MS, kDa | MP, kDa (%) | SAXS, kDa | SEC, kDa | Swiss model calc. | Uniprot (kDa) and oligomerization | PDB | Protein abundance (copy number per cell) | |
|---|---|---|---|---|---|---|---|---|---|---|
| concentrations | 1.875 - 40 µM | | 50nM | 7-60 µM<br>0.1-0.6mg\ml | 13.32 µM | | | 2EHJ | Ref.1 | Ref.2 |
| Monomer | | | | | | | 22.53 | | 2780 | 4260 |
| Dimer | | 44.95 | 44 (61%) | | 52 (40-66) | | Homodimer or homotrimer in the absence of substrates, and homopentamer or homohexamer in the presence of substrates. | | highly expressed protein | |
| Trimer | | | | | | | | | | |
| Tetramer | | 89.89 | 89 (13%) | *I(0)* = 101.4 (σ=2.9) | 88 (69-113) | Homotetramer | | | | |
| Hexamer | | | 131 (9%) | | 130 (101-166) | | | | | |
| Octamer | | | 175 (5%) | | | | | | | |
| Graph | 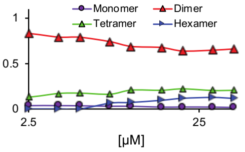 | | 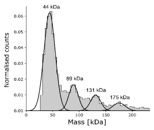 | | 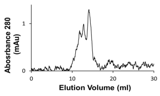 | | | | | |





#### SpeB- P60651

| Method | MS conc. | MS, kDa | MP, kDa (%) | SAXS, kDa | SEC, kDa | Swiss model calc. | Uniprot (kDa) and oligomerization | PDB | Protein abundance (copy number per cell) | |
|---|---|---|---|---|---|---|---|---|---|---|
| concentrations | 7.5 - 40 µM | | 110 nM | 8-63 µM<br>0.3-2 mg\ml | 9 µM | | | 7LBA | Ref.1 | Ref.2 |
| Monomer | | | | | | | 33.56 | | 3530 | 1063 |
| Dimer | | | 42 (25%) | | | | | | highly expressed protein | |
| Trimer | | 100.54 | 105 (33%) | | 110 (86-140) | | | | | |
| Tetramer | | | | | | | | | | |
| Hexamer | | 201.08 | 206 (15%) | *I(0)* = 175.5 (σ=7.3) | 203 (159-260) | Hexamer | | | | |
| Graph | 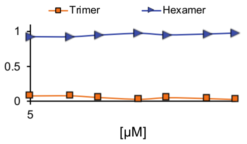 | | 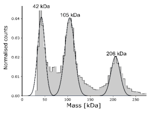 | | 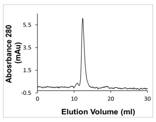 | | | | | |





#### IspD- Q46893

| Method | MS conc. | MS, kDa | MP, kDa (%) | SAXS, kDa | SEC, kDa | Swiss model calc. | Uniprot (kDa) and oligomerization | PDB | Protein abundance (copy number per cell) | |
|---|---|---|---|---|---|---|---|---|---|---|
| concentrations | 7.5 - 40 µM | | NA<br>Under the detection limit of MP | 9-72 µM<br>0.23-1.9 mg\ml | 12 µM | | | 1VGT, 3N9W, 1i52 | Ref.1 | Ref.2 |
| Monomer | | | | | 43 (34-56) | | 25.61 | | low expressed proteins | |
| Dimer | | 51.35 | | | | Homodimer | Homodimer | | | |
| Trimer | | | | *I(0)* = 99.6 (σ=3.5) | 84 (66-108) | | | | | |
| Tetramer | | 102.71 | | | 133 (104-170) | | | | | |
| Hexamer | | | | | | | | | | |
| Graph | 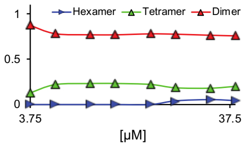 | | | | 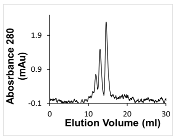 | | | | | |




#### BaeR- P69228

| Method | MS conc. | MS, kDa | MP, kDa (%) | SAXS, kDa | SEC, kDa | Swiss model calc. | Uniprot (kDa) and oligomerization | PDB | Protein abundance (copy number per cell) | |
|---|---|---|---|---|---|---|---|---|---|---|
| concentrations | 0.625 - 40 µM | | NA<br>Under the detection limit of MP | 13-107 µM<br>0.4-2.95 mg\ml | 11 µM | | | 4B09 | Ref.1 | Ref.2 |
| Monomer | | 27.60 | | *I(0)* = 22.9 (σ=0.9) | 25 (20-32) | | 27.66 | | low expressed proteins | |
| Dimer | | | | | | dimer | dimer | | | |
| Graph | 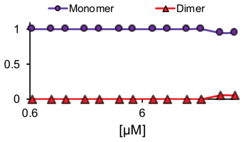 | | | | 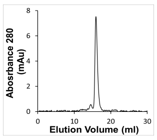 | | | | | |

#### AcuI P26646

| Method | MS conc. | MS, kDa | MP, kDa (%) | SAXS, kDa | SEC, kDa | Swiss model calc. | Uniprot (kDa) and oligomerization | PDB | Protein abundance (copy number per cell) | |
|---|---|---|---|---|---|---|---|---|---|---|
| concentrations | 0.625 - 40 µM | | 45 nM | 4-29µM<br>0.125-1 mg\ml | 134 nM | | | | Ref.1 | Ref.2 |
| Monomer | | 34.68 | 39 (35%)* fitted: 38(49%) | | 56 (44-72) | | 34.73 | | 186 | 882 |
| Dimer | | 69.36 | 69 (47%) fitted: 67 (46%) | *I(0)* =209.8 (σ=11.4) | | Homodimer | Homodimer | | | |
| Trimer | | | | | | | | | | |
| Tetramer | | 104.04 | 134 (4%) fitted: 108(5%) | | 126 (98-161) | | | | | |
| Hexamer | | | | | 201 (157-258) | | | | | |
| Graph | 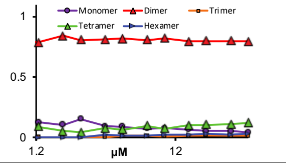 | | 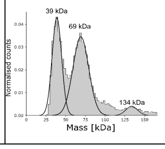 | | 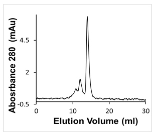 | | | 1O89<br>108C | low expressed proteins | |







#### PyrF P08244

| Method | MS conc. | MS, kDa | MP, kDa (%)*** | SAXS, kDa | SEC, kDa | Swiss model calc. | Uniprot (kDa) and oligomerization | PDB | Protein abundance (copy number per cell) | |
|---|---|---|---|---|---|---|---|---|---|---|
| concentrations | 0.234- 40 µM | | 60 nM | 10-81µM<br>0.3-2 mg\ml | 11 µM | | | | Ref.1 | Ref.2 |
| Monomer | | 26.31 | | | 44 (35-57) | | 26.35 | | 212 | 309 |
| Dimer | | 52.61 | 58 (66%) | *I(0)* = 57 (σ=4.1) | | Homodimer | Homodimer | | | |
| Trimer | | | | | | | | | | |
| Graph | 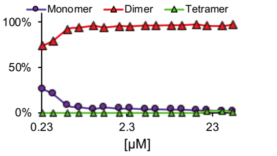 | | 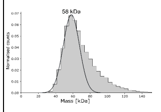 | | 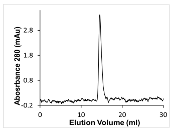 | | | 1EIX, 1L2U | low expressed proteins | |







#### ThiD P76422

| Method | MS conc. | MS, kDa | MP, kDa (%) | SAXS, kDa | SEC, kDa | Swiss model calc. | Uniprot (kDa) and oligomerization | PDB | Protein abundance (copy number per cell) | |
|---|---|---|---|---|---|---|---|---|---|---|
| concentrations | 5 - 40 µM | | 50 nM | | 10.5 µM | | | | Ref.1 | Ref.2 |
| Monomer | | 28.61 | | | 41 (32-52) | | 28.64 | | 186 | |
| Dimer | | 57.21 | 58 (81%) | | | Homodimer | Monomer | | | |
| Graph | 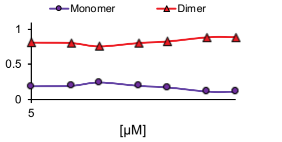 | | 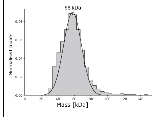 | | 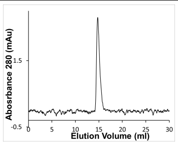 | | | NA | low expressed proteins | |







#### NadE- P18843

| Method | MS conc. | MS, kDa | MP, kDa (%) | SAXS, kDa | SEC, kDa | Swiss model calc. | Uniprot (kDa) and oligomerization | PDB | Protein abundance (copy number per cell) | |
|---|---|---|---|---|---|---|---|---|---|---|
| concentrations | 0.938 - 40 µM | | 66 nM | 9-74 µM<br>0.2-1.7 mg\ml | 11 µM | | | | Ref.1 | Ref.2 |
| Monomer | | | | | | | 27.16 | | 746 | 598 |
| Dimer | | 61.21 | 66 (83%) | *I(0)* = 90.5 (σ=5.8) | 46 (36-59) | Homodimer | Homodimer | | | |
| Tetramer | | | | | | | | | | |
| Graph | 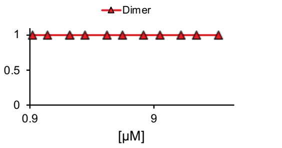 | | 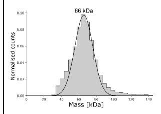 | | 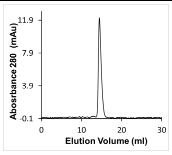 | | | 1WXF, 1WXI | low expressed proteins | |







#### Crp P0ACJ8

| Method | MS conc. | MS, kDa | MP, kDa (%) | SAXS, kDa | SEC, kDa | Swiss model calc. | Uniprot (kDa) and oligomerization | PDB | Protein abundance (copy number per cell) | |
|---|---|---|---|---|---|---|---|---|---|---|
| concentrations | 0.938 - 40 µM | | 48 nM | | 13 µM | | | | Ref.1 | Ref.2 |
| Monomer | | | | | 31 (24-40) | Monomer, Homodimer | 23.64 | | 1980 | 3463 |
| Dimer | | 47.16 | 48 (94%) | | | | Dimer | | | |
| Graph | 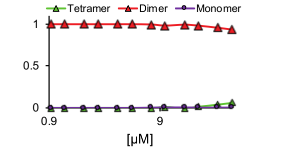 | | 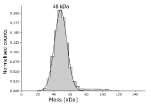 | NA | 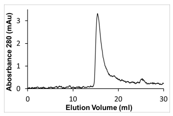 | | | 2GZW, 3N4M, 5CIZ,1LB2 | low expressed proteins | |








**Ref.1** Ishihama, Y., Schmidt, T., Rappsilber, J., Mann, M., Hartl, F. U., Kerner, M. J., & Frishman, D. (2008). Protein abundance profiling of the Escherichia coli cytosol. BMC Genomics, 9(1), 102.

**Ref.2** Fauvet, Bruno, et al. "Bacterial Hsp90 mediates the degradation of aggregation-prone Hsp70-Hsp40 substrates preferentially by HslUV proteolysis," bioRxiv (2018): 451989.


emPAI-derived copy no/cell-

*Calculated using 1fL as the volume of the cell, protein concentration*avogadro no. * cell volume
