## Supplementary Table S3 for "Protein Quaternary Structures in Solution are a Mixture of Multiple forms"

| Supplementary Table S3- Alpha Fold results of all 17 E.coli protiens | | | | | |
|---|---|---|---|---|---|
| **Oligomeric state** | **Sample name** | **pTM** | **plDDT** | **ipTM** | **PAE** |
| Monomer | 2_NfuA | 0.54 | 84.5 | | 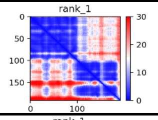 |
| Dimer | 2_NfuA_dimer | 0.4 | | 0.18 | 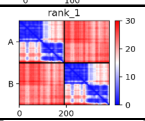 |
| Trimer | 2_NfuA_trimer | 0.34 | | 0.16 | 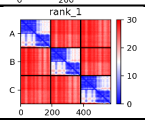 |
| Tetramer | 2_NfuA_tetramer | 0.33 | | 0.21 | 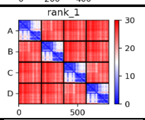 |
| Hexamer | 2_NfuA_hexamer | 0.27 | | 0.2 | 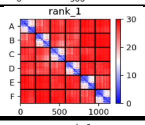 |
| Monomer | 3_NadK | 0.83 | 91.4 | | 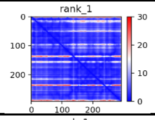 |
| Dimer | 3_NadK_dimer | 0.92 | | 0.91 | 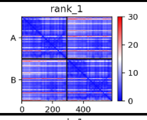 |
| Trimer | 3_NadK_trimer | 0.8 | | 0.75 | 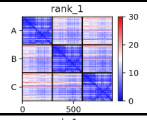 |
| Tetramer | 3_NadK_tetramer | 0.89 | | 0.88 | 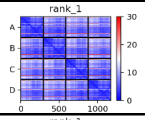 |
| Hexamer | 3_NadK_hexamer | 0.39 | | 0.33 | 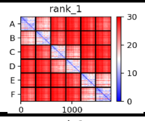 |
| Monomer | 4_FabG | 0.87 | 97 | | 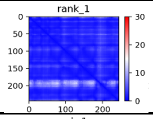 |
| Dimer | 4_FabG_dimer | 0.95 | | 0.94 | 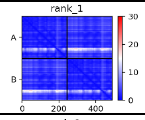 |
| Trimer | 4_FabG_trimer | 0.81 | | 0.77 | 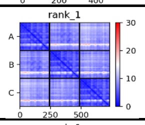 |
| Tetramer | 4_FabG_tetramer | 0.94 | | 0.93 | 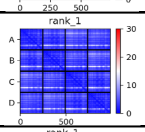 |
| Hexamer | 4_FabG_hexamer | 0.78 | | 0.76 | 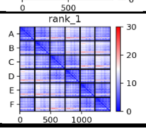 |

| | | | | | |
|---|---|---|---|---|---|
| Monomer | 5_DeoC | 0.85 | 96.1 | | 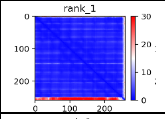 |
| Dimer | 5_DeoC_dimer | 0.93 | | 0.92 | 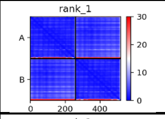 |
| Trimer | 5_DeoC_trimer | 0.6 | | 0.45 | 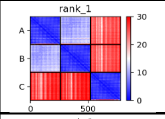 |
| Tetramer | 5_DeoC_tetramer | 0.53 | | 0.41 | 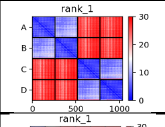 |
| Hexamer | 5_DeoC_hexamer | 0.37 | | 0.27 | 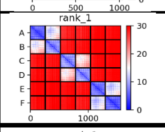 |
| Monomer | 6_eftS | 0.8 | 94.2 | | 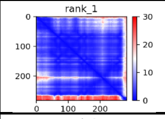 |
| Dimer | 6_eftS_dimer | 0.61 | | 0.42 | 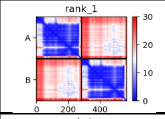 |
| Trimer | 6_eftS_trimer | 0.41 | | 0.2 | 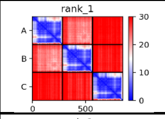 |
| Tetramer | 6_eftS_tetramer | 0.42 | | 0.28 | 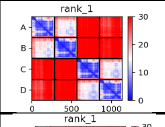 |
| Hexamer | 6_eftS_hexamer | 0.34 | | 0.25 | 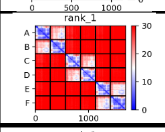 |
| Monomer | 7_SodA | 0.86 | 97.8 | | 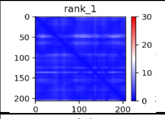 |
| Dimer | 7_SodA_dimer | 0.95 | | 0.93 | 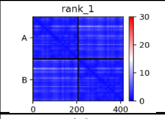 |
| Trimer | 7_SodA_trimer | 0.66 | | 0.51 | 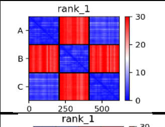 |
| Tetramer | 7_SodA_tetramer | 0.52 | | 0.37 | 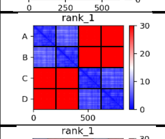 |
| Hexamer | 7_SodA_hexamer | 0.39 | | 0.29 | 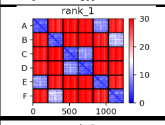 |
| Monomer | 9_gpmA | 0.85 | 95.7 | | 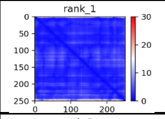 |
| Dimer | 9_gpmA_dimer | 0.95 | | 0.94 | 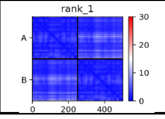 |

| | | | | | |
|---|---|---|---|---|---|
| Trimer | 9_gpmA_trimer | 0.54 | | 0.35 | 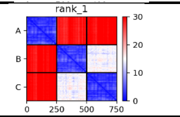 |
| Tetramer | 9_gpmA_tetramer | 0.52 | | 0.37 | 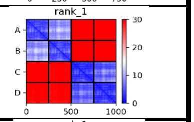 |
| Hexamer | 9_gpmA_hexamer | 0.39 | | 0.3 | 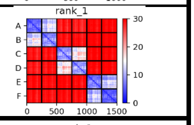 |
| Monomer | 11__can | 0.84 | 95.3 | | 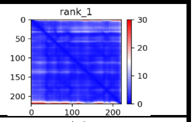 |
| Dimer | 11__can_dimer | 0.94 | | 0.94 | 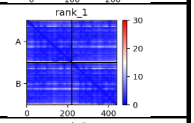 |
| Trimer | 11__can_trimer | 0.9 | | 0.87 | 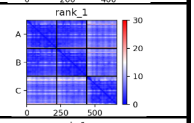 |
| Tetramer | 11__can_tetramer | 0.95 | | 0.94 | 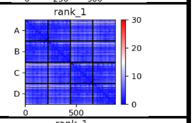 |
| Hexamer | 11__can_hexamer | 0.44 | | 0.36 | 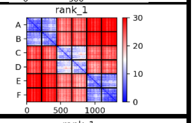 |
| Monomer | 12__upp | 0.85 | 95.8 | | 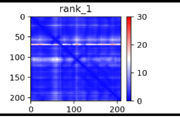 |
| Dimer | 12__upp_dimer | 0.94 | | 0.93 | 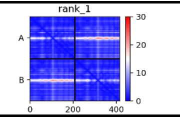 |
| Trimer | 12__upp_trimer | 0.72 | | 0.67 | 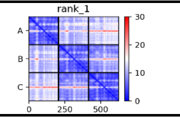 |
| Tetramer | 12__upp_tetramer | 0.92 | | 0.91 | 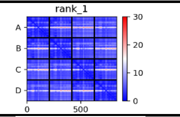 |
| Hexamer | 12__upp_hexamer | 0.58 | | 0.53 | 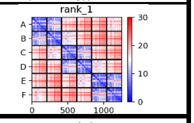 |
| Monomer | 13_speB | 0.87 | 95.5 | | 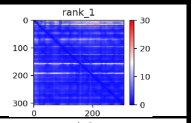 |
| Dimer | 13_speB_dimer | 0.73 | | 0.53 | 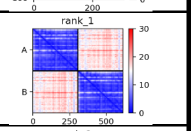 |
| Trimer | 13_speB_trimer | 0.86 | | 0.82 | 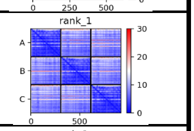 |
| Tetramer | 13_speB_tetramer | 0.66 | | 0.58 | 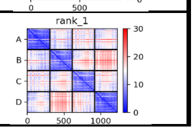 |

| | | | | | |
|---|---|---|---|---|---|
| Hexamer | 13_speB_hexamer | 0.87 | | 0.86 | 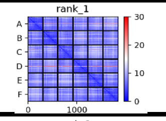 |
| Monomer | 14_ispD | 0.83 | 92.5 | | 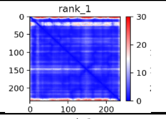 |
| Dimer | 14_ispD_dimer | 0.89 | | 0.9 | 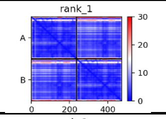 |
| Trimer | 14_ispD_trimer | 0.56 | | 0.43 | 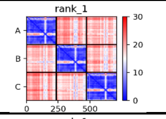 |
| Tetramer | 14_ispD_tetramer | 0.48 | | 0.38 | 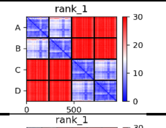 |
| Hexamer | 14_ispD_hexamer | 0.36 | | 0.29 | 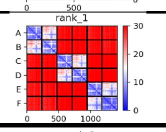 |
| Monomer | 15_BaeR | 0.54 | 79.1 | | 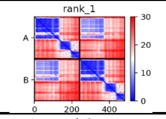 |
| Dimer | 15_BaeR_dimer | 0.54 | | 0.48 | 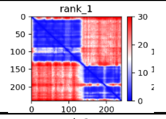 |
| Trimer | 15_BaeR_trimer | 0.39 | | 0.3 | 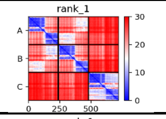 |
| Tetramer | 15_BaeR_tetramer | 0.34 | | 0.24 | 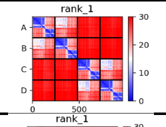 |
| Hexamer | 15_BaeR_hexamer | 0.28 | | 0.21 | 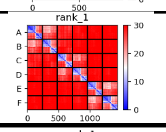 |
| Monomer | 16_AcuI | 0.87 | 96.2 | | 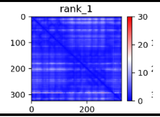 |
| Dimer | 16_AcuI_dimer | 0.94 | | 0.94 | 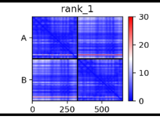 |
| Trimer | 16_AcuI_trimer | 0.47 | | 0.31 | 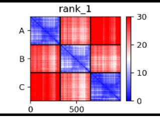 |
| Tetramer | 16_AcuI_tetramer | 0.5 | | 0.37 | 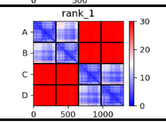 |
| Hexamer | 16_AcuI_hexamer | 0.38 | | 0.3 | 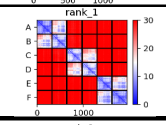 |
| Monomer | 17_pyrF | 0.84 | 93.8 | | 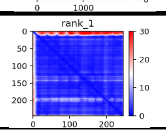 |

| | | | | | |
|---|---|---|---|---|---|
| Dimer | 17_pyrF_dimer | 0.92 | | 0.91 | 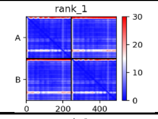 |
| Trimer | 17_pyrF_trimer | 0.5 | | 0.34 | 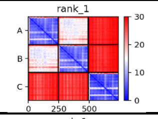 |
| Tetramer | 17_pyrF_tetramer | 0.51 | | 0.37 | 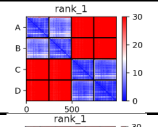 |
| Hexamer | 17_pyrF_hexamer | 0.38 | | 0.29 | 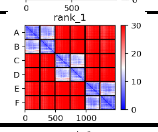 |
| Monomer | 19_thiD | 0.86 | 93.7 | | 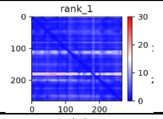 |
| Dimer | 19_thiD_dimer | 0.95 | | 0.94 | 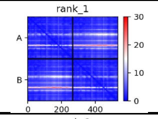 |
| Trimer | 19_thiD_trimer | 0.52 | | 0.36 | 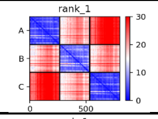 |
| Tetramer | 19_thiD_tetramer | 0.52 | | 0.41 | 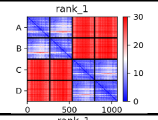 |
| Hexamer | 19_thiD_hexamer | 0.4 | | 0.31 | 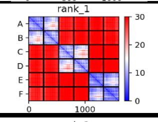 |
| Monomer | 23_nadE | 0.86 | 95.4 | | 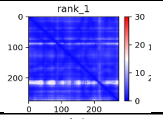 |
| Dimer | 23_nadE_dimer | 0.95 | | 0.95 | 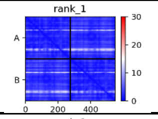 |
| Trimer | 23_nadE_trimer | 0.54 | | 0.41 | 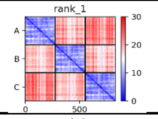 |
| Tetramer | 23_nadE_tetramer | 0.52 | | 0.39 | 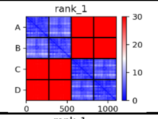 |
| Hexamer | 23_nadE_hexamer | 0.4 | | 0.3 | 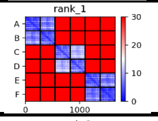 |
| Monomer | 27_crp | 0.8 | 94.4 | | 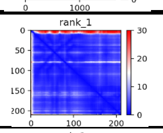 |
| Dimer | 27_crp_dimer | 0.92 | | 0.92 | 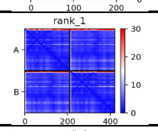 |
| Trimer | 27_crp_trimer | 0.83 | | 0.81 | 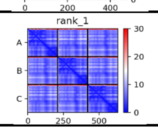 |

| | | | | | |
|---|---|---|---|---|---|
| Tetramer | 27_crp_tetramer | 0.5 | | 0.37 | 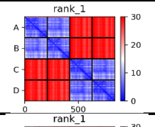  |
| Hexamer | 27_crp_hexamer | 0.38 | | 0.28 | 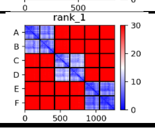  |
